# Meiotic gene transcription programs are mediated by A-MYB and BRDT-dependent RNA polymerase II pause release during mammalian prophase I

**DOI:** 10.1101/2022.08.19.504615

**Authors:** Adriana K. Alexander, Edward J. Rice, Gilad Barshad, Lina Zhu, Paula E Cohen, Charles G. Danko

## Abstract

During meiotic prophase I, germ cells must balance transcriptional activation with meiotic recombination and chromosome synapsis, biological processes requiring extensive changes to chromatin state and structure. Here we explored the interplay between chromatin accessibility and transcription across a detailed time-course of murine male meiosis by measuring genome-wide patterns of chromatin accessibility, nascent transcription, and processed mRNA. To understand the relationship between these parameters of gene regulation and recombination, we integrated these data with maps of double-strand break formation. Maps of nascent transcription show that Pol II is loaded on chromatin and maintained in a paused state early during prophase I. In later stages of prophase I, paused Pol II is released in a coordinated transcriptional burst resulting in ∼3-fold increase in transcription. Release from pause is mediated by the transcription factor A-MYB and the testis-specific bromodomain protein, BRDT. The burst of transcriptional activity is both temporally and spatially segregated from key steps of meiotic recombination: double strand breaks show evidence of chromatin accessibility earlier during prophase I and at distinct loci from those undergoing transcriptional activation, despite shared chromatin marks. Our findings reveal the mechanism underlying chromatin specialization in either transcription or recombination in meiotic cells.

## INTRODUCTION

Spermatogenesis is a multistep process required for the daily production of millions of spermatozoa within the seminiferous epithelium of the testis^1^. Meiosis is a critical stage in spermatogenesis during which the genome is first replicated and then halved to produce four haploid gametes. The first and longest stage of meiosis is prophase I, during which dramatic changes in chromatin occur to facilitate the events of synapsis and meiotic recombination^2^. Upon entry into prophase I, the topoisomerase-like protein SPO11, together with multiple accessory proteins, catalyzes the induction of hundreds of DNA double-strand breaks (DSBs) across the genome^3^. Of these, 90% will resolve as non-crossovers, while the remaining 10% will resolve as crossovers, which result in the physical connection between two distinct chromosomes^4^. In mammals, DSB induction initiates the assembly of a tripartite protein structure called the synaptonemal complex (SC), which tethers homologous chromosomes and facilitates the progressive repair of DSBs^5,6^. By the end of prophase I, when the SC begins to break down, the DSB-initiated crossovers tether homologous chromosomes until the first meiotic division ^2,7^. These events, from DSB formation through the first meiotic division, require a complicated and intricate series of changes to chromatin that are essential for both meiotic recombination and the proper segregation of DNA into haploid gametes.

In addition to meiotic recombination, prophase I spermatocytes have an essential role in enacting complex transcriptional programs that control events during meiosis as well as the final differentiation stages of spermatogenesis, known as spermiogenesis^8^. Prophase I spermatocytes initiate the transcription of thousands of protein-coding genes, pseudogenes, piRNAs, long non-coding RNAs, and transposable elements, establishing a highly complex transcriptional program^8–13^. Many of the genes transcribed during this stage are necessary to facilitate meiotic recombination and synapsis during prophase I itself^14^. Yet, RNAs transcribed during prophase I may also be stored until they are needed to facilitate events during transcriptionally inert stages later in spermiogenesis^13,15^. Dysfunctional gene expression during prophase I can lead to a variety of infertility phenotypes that are characterized by either meiotic arrest or spermiogenic failure^10,16–19^.

Relatively little is known about how transcription programs are controlled during prophase I. The sequence-specific transcription factor A-MYB is known to activate the transcription of protein-coding genes and piRNAs by binding to proximal and distal regulatory elements and super enhancers across the genome^19–22^. The A-MYB protein, encoded by the *Mybl1* gene, first appears in the pachytene stage of prophase I^19^. Once expressed, A-MYB transforms the regulatory environment by switching on a large number of enhancers that were inactive during earlier stages of spermatogenesis^21^. However, it remains unclear how A-MYB activates transcription during prophase I. First, we know little about how A-MYB identifies target enhancers carrying its information-poor consensus sequence, and, beyond a number of well-characterized piRNAs, the identity of its direct target genes^20^. Second, we do not know the mechanistic basis by which A-MYB activates transcription of its target genes. Finally, and perhaps most importantly, a general feature of transcriptional activation is extensive decondensation of chromatin. Therefore, a central question is how spermatocytes are able to balance the separate, and arguably opposing, tasks of chromatin decondensation to facilitate transcription and chromosome condensation to aid the defining events in prophase I.

Here we integrated genomic data throughout meiosis to ask how spermatocytes achieve a balance between the hallmark events of prophase I and transcriptional activation. We performed the first comprehensive analysis of gene expression and chromatin accessibility at discrete substages of meiotic prophase I: leptonema, zygonema, pachynema, and diplonema. To understand the mechanistic basis for the dynamic changes in transcriptional activity during meiotic recombination, we studied nascent transcription by mapping the location of transcriptionally engaged RNA polymerase II (Pol II) using length-extension chromatin run-on and sequencing (leChRO-seq) to avoid the stability-derived biases of steady-state mRNA sequencing^23^. Our stage-resolved maps of nascent transcription show that Pol II is loaded on chromatin and maintained in a paused state during early prophase I. Starting in pachynema, paused Pol II is released into productive elongation in a global and highly coordinated burst of transcriptional activity during which transcription is increased by ∼3-fold over baseline levels. Transcriptional activation during pachynema is initiated by A-MYB, which recruits the testis-specific bromodomain protein, BRDT, to release paused Pol II into productive elongation. Both the chromatin environment and paused Pol II, which allow A-MYB binding during pachynema to activate transcription, are established by the activity of other pioneer transcription factors during earlier stages of meiosis. Finally, sites of transcriptional activity are both temporally and spatially segregated from those involved in meiotic DSB repair and recombination despite a largely shared epigenetic environment characterized by high H3K4me3 levels, allowing cells to activate transcription of essential genes and undergo DSB repair at distinct genomic loci. Taken together, these findings reveal a separation of chromatin domains, in which distinct loci are programmed during early prophase I to focus on either meiotic recombination or transcriptional activation.

## RESULTS

### Transcriptionally active RNA Polymerase II is enriched in pachynema

Prophase I can be divided into 5 substages defined by the status of the SC: formation of the SC begins in leptonema, becomes progressively connected in zygonema, fully synapsed in pachynema, and disassembles during diplonema and diakinesis^2^. Meiotic recombination occurs alongside these events, with DSBs initiated in leptonema, homolog pairing, DSB repair and the appearance of non-crossovers in zygonema, and crossovers arising in pachynema. To investigate the pattern of transcriptional activity through successive substages of prophase I, we measured the abundance of RNA polymerase II (Pol II) on chromatin by immunofluorescence on chromosome spreads using antibodies that recognize the N-terminal region of the largest Pol II subunit, RPB1 (**Fig. 1A-C**). Immunofluorescent imaging of prophase I substages showed significant enrichment (>3-fold-increase) of chromatin-associated Pol II in pachynema and diplonema compared to leptonema and zygonema (n_mice_ = 3; n_cells_ = 606; \*\*\*\**p*-value < 0.0001; **Fig. 1 A and D**). Pachynema and diplonema showed diffuse Pol II signal throughout the nucleus, except in regions of pericentromeric heterochromatin (**Fig. S1**, arrowheads). Pol II signal was also markedly reduced within the sex body (SB), the heterochromatin-rich sub-domain of the nucleus in which the XY bivalent resides, in pachynema and diplonema^24^. These data show that chromatin-associated Pol II increases from zygonema to pachynema, predominantly in euchromatic regions on the autosomes.

**Figure 1.**
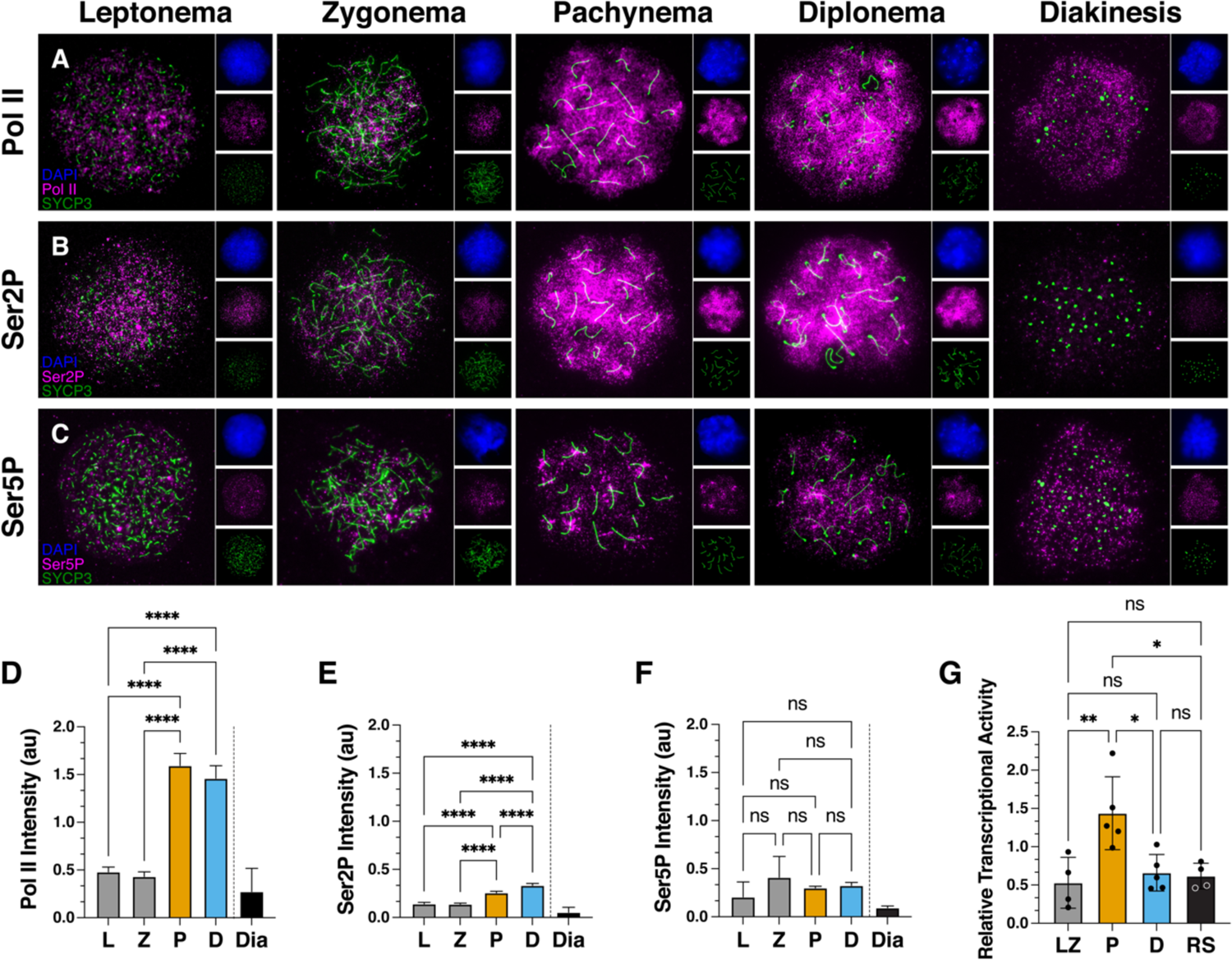
Transcriptionally active isoform of RNA Polymerase II, Ser2P, is enriched in pachynema and diplonema. A-C) Immunofluorescence staining on spread meiotic spermatocyte preparations from wildtype C57Bl/6 mice using antibodies against SYCP3 (green) and Pol II (A), Ser2P (B), and Ser5P (C) (magenta). Leptonema (L), zygonema (Z), pachynema (P), diplonema (D), and diakinesis (Dia) staged spermatocytes are defined by SYCP3 morphology. D-F) Quantification of signal intensity for Pol II, Ser2P, and Ser5P immunofluorescence staining in wildtype spermatocytes. Average intensity is shown, and error bars represent 95% confidence intervals. Samples were obtained from three 3-month-old mice. D) *n* = 606 prophase I cells. E) *n* = 677 prophase I cells. F) There was no significant difference in the fluorescence intensity of Ser5P between prophase I substages. *n* = 586 prophase I cells. G) Radioactive nuclear run-on results comparing the relative levels of nascent transcription between leptonema/zygonema (LZ), pachynema (P), diplonema (D), and round spermatids (RS). Average relative transcriptional activity is shown, and error bars represent standard deviation. D-G) \**p*-value < 0.0117, \*\**p*-value < 0.0056, **** *p*-value < 0.0001, One-way ANOVA with the post-hoc Tukey’s multiple comparison’s test.

To connect chromatin-associated Pol II with transcriptional output, we next examined the transcriptional state of the Pol II holoenzyme during each stage of prophase I. Different stages of the Pol II transcription cycle are associated with differences in the phosphorylation states of the C-terminal domain (CTD) of RPB1^25^. We performed immunofluorescence of chromosome spreads using antibodies recognizing phosphorylation of Ser2 (Ser2P), which marks the elongating form of Pol II, and Ser5P, which is associated with Pol II in a promoter-proximal paused state^25^ (**Fig. 1 B-C, E-F**). We found significantly higher levels of total Ser2P signal in pachynema and diplonema when compared to leptonema and zygonema (n_mice_ = 3; n_cells_ = 677; \*\*\*\**p*-value < 0.0001; **Fig. 1 B and E**). However, there was no significant difference in Ser5P signal intensity across prophase I substages even though total Pol II was detected at low intensity in leptonema and zygonema (n_mice_ = 3; n_cells_ = 586; *p*-value > 0.05; **Fig. 1 B and E**). Taken together, these results suggest that the Pol II complexes observed in pachynema and diplonema are primarily transcriptionally active, whereas those in leptonema and zygonema are enriched for a paused transcriptional state.

To measure the amount of transcriptionally engaged RNA polymerase (Pol) in a more quantitative manner, we next used nuclear run-on assays to analyze each stage of prophase I and early stages during spermiogenesis. Nuclear run-ons use the activity of transcriptionally engaged Pol I-III to incorporate radioactive [α32P]CTP, which is measured using a scintillation counter. Notably, this approach has two advantages compared with immunofluorescence: first, it directly measures transcriptionally engaged RNA polymerases, and second it provides a quantitative readout of RNA polymerase abundance that is linear over many orders of magnitude^26,27^. Transcriptional activity was nearly three times greater (2.7 fold increase) in pachynema than in leptonema/zygonema, diplonema, and round spermatids (n = 21; \*\**p*-value = 0.0029; **Fig. 1G**). In contrast to immunofluorescence measurements, radioactive counts were not significantly higher in diplonema compared with leptonema/zygonema, which may be explained either by chromatin-bound RNA polymerase that was not transcriptionally engaged, substantially decreased Pol I or III in diplonema, or technical differences between experiments. Taken together, our results demonstrate the existence of a sharp burst of transcriptional activity that peaks during pachynema, perhaps by releasing paused Pol II into an active state.

**Figure S1.**
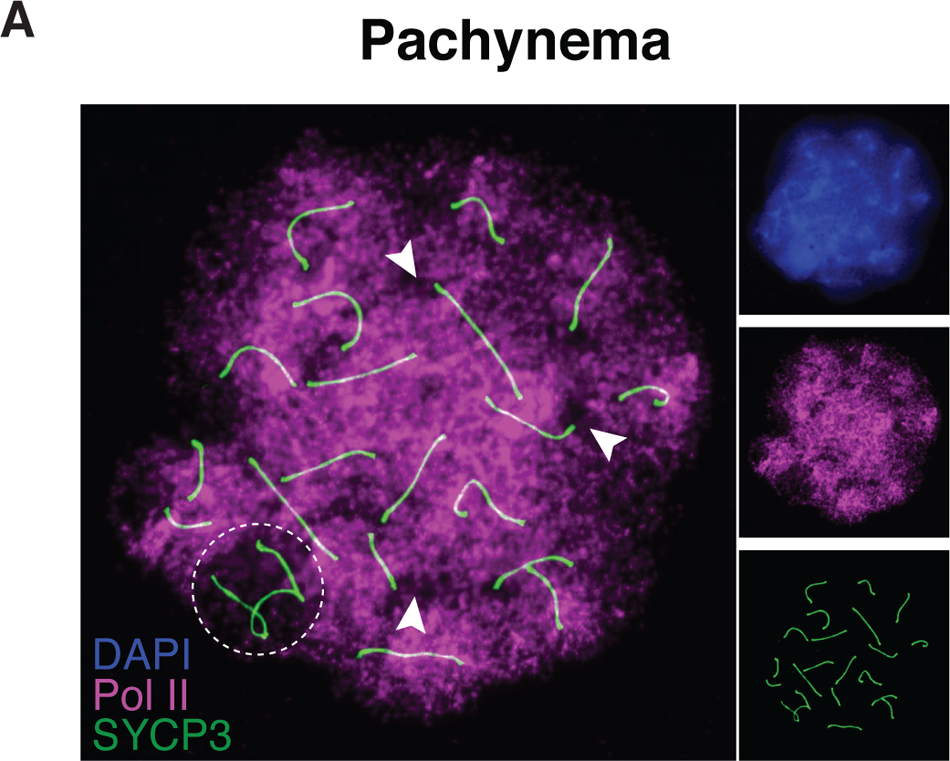
Localization of RNA Pol II on a pachytene spermatocyte chromosome spread. A) Immunolocalization of Pol II (magenta), SYCP3 (green), and DAPI (blue) in WT pachytene spermatocytes. The sex body, housing the X and Y chromosomes, is outlined in white. Arrows indicate absence of Pol II signal near pericentromeric heterochromatin.

### Transcriptional activation in pachynema is driven by pause release

To determine how the robust burst of transcriptional activation observed during prophase I affects the status of transcription and chromatin on individual genes, we next measured genome-wide patterns of transcription and chromatin accessibility. We isolated cells representing leptotene/zygotene, pachytene, and diplotene stages of prophase I using either STA-PUT or FACS (see Methods). Staining against proteins that are defining markers of different prophase I substages (SYCP3 and γH2AX) supported a high purity in each fraction (83.5%, 77.5%, and 87%, for leptonema/zygonema, pachynema, and diplonema, respectively), with the majority of contamination representing elongated spermatids in all fractions. We analyzed each cell stage using three genome-wide molecular assays that measure different layers of gene regulation (**Fig. 2A**): ATAC-seq to measure chromatin accessibility, leChRO-seq to map the location and orientation of RNA Pol I-III, and RNA-seq to measure the abundance of processed mRNA. This combination of molecular assays allowed us to identify open chromatin, classify promoter and enhancer elements based on the transcription of enhancer-templated RNAs (eRNAs)^28^, and measure gene expression changes at the level of both transcription and steady-state mRNA abundance. To adjust for global changes in transcription observed during prophase I, we used radioactive nuclear run-on assays to normalize leChRO-seq signal. After sequencing 2-4 replicates from each molecular assay to a depth of 16-87 million uniquely mapped reads, we verified that the majority of variation in each sequencing library represented the stage during prophase I (PC1: 77%, 93%, and 95% for leChRO-seq, RNA-seq, and ATAC-seq, respectively) (**Fig. 2B**).

**Figure 2.**
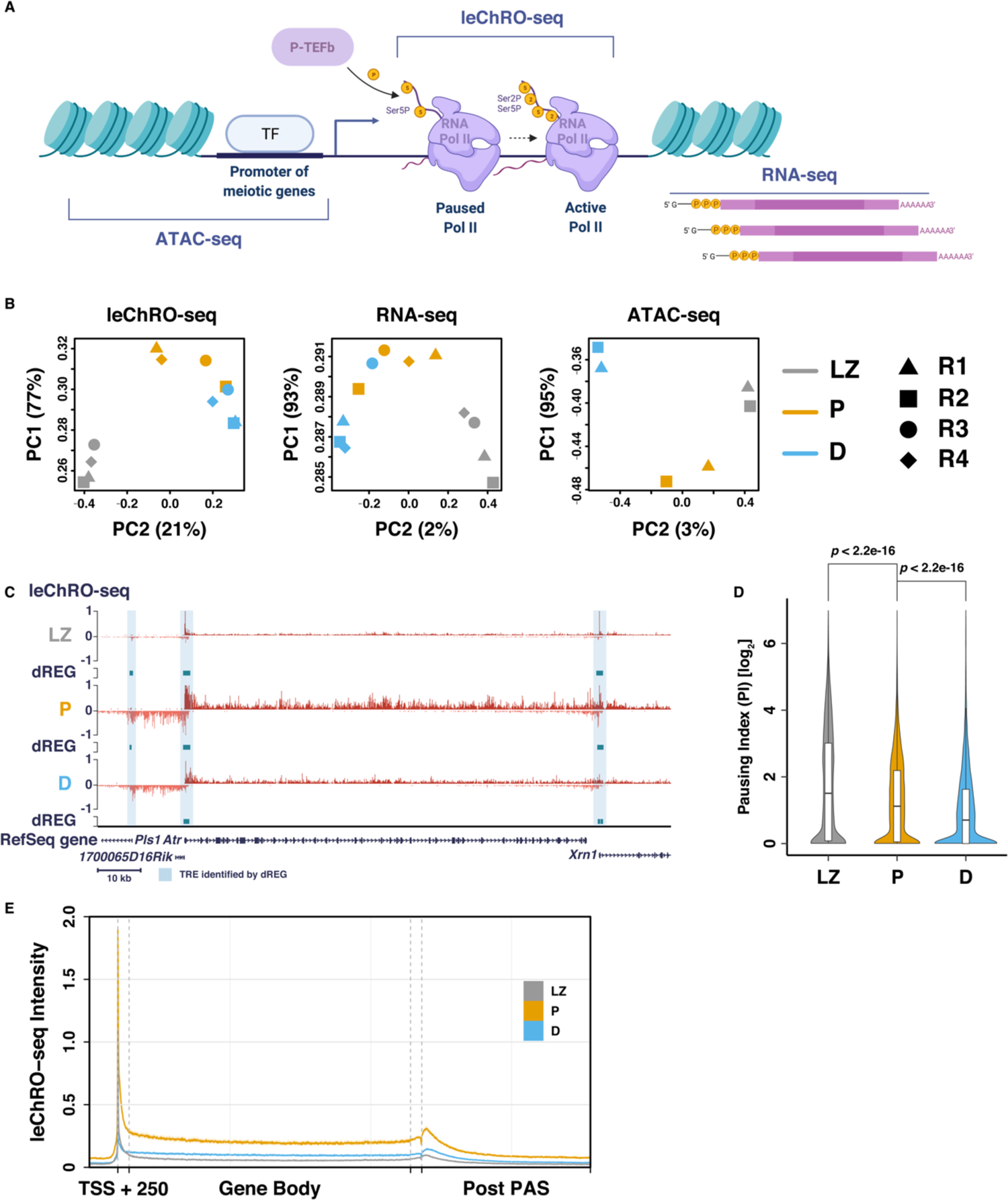
leChRO-seq detects nascent transcription in prophase I spermatocytes. A) Schematic of key steps in the transcription cycle. The transcription start site (TSS) is labeled with an arrow; nucleosomes are depicted in teal; nascent RNA is shown in dark red; and mRNA is shaded in dark pink. ATAC-seq detects accessible regulatory elements, such as promoters and enhancers. leChRO-seq provides base pair resolution of engaged and active Pol II across the genome. RNA-seq profiles poly-adenylated mRNA transcripts. B) Principal component analysis (PCA) plots showing variation between replicate sets of prophase I substages for leChRO-seq, RNA-seq, and ATAC-seq libraries. C) leChRO-seq signal at the *Atr* locus for LZ (top), P (center), and D (bottom). dREG peaks (teal) are shown for each prophase I substage. TSSs overlapping with dREG peak calls are shaded in blue. D) Violin plots of pause index values calculated from the ratio of pause peak density to gene body density of leChRO-seq reads. *p* < 2.2e-16, Wilcoxon matched-pairs signed rank test. E) Metagene plot of median leChRO-seq signal intensity at annotated gene boundaries. LZ: leptonema/zygonema; pachynema: P; diplonema: D.

Immunofluorescence showed an accumulation of Ser5P throughout prophase I (**Fig. 1C**), which may reflect Pol II in a promoter-proximal paused state. To confirm this finding, we first focused our analysis on leChRO-seq and asked whether Pol II accumulated near the promoter in a promoter-proximal pause position, classically ∼20-60 bp downstream of the transcription start site (TSS). Consistent with this model, examination of leChRO-seq data revealed an accumulation of promoter-proximal Pol II in leptonema/zygonema. For example, *Atr, Xrn1*, and *1700065D16Rik* all had high levels of Pol II within 60 bp of their TSS in leptonema/zygonema (*Atr*: Ataxia telangiectasia and Rad3 related, required for homologous recombination and synapsis^29^; *Xrn1*: 5’-3’ exoribonuclease 1, involved in replication-dependent histone mRNA degradation^30^; *1700065D16Rik*: uncharacterized protein, expression restricted to the testis^31^). Progression through pachynema and diplonema was associated with increased Pol II in the gene-body of all three genes (**Fig. 2C**).

To quantitatively examine the pattern of paused Pol II genome-wide, we computed the pausing index, or the density of paused Pol II signal relative to the signal across the gene body, which can be interpreted as the rate of pause release^32^, during each stage of spermatogenesis (**Fig. 2D-E**). Pausing index was highest in leptonema/ zygonema, and decreased during successive stages of prophase I. By contrast, gene body transcription increases as prophase I proceeds, peaking in pachynema and dropping slightly in diplonema. We conclude that increased transcriptional activity in pachytene cells is driven by paused Pol II being released into productive elongation after first being established during leptonema/zygonema.

### Pachytene transcriptional burst activates genes involved in meiosis, transcription, and spermatogenesis

We next analyzed which annotated protein-coding genes, pseudogenes, and non-coding RNAs showed transcriptional changes during the transition between leptonema/zygonema, pachynema, and diplonema. Differential expression analysis of leChRO-seq data identified 9,062 protein-coding genes, 103 pseudogenes, and 389 long non-coding RNAs (lncRNAs) that were either up- or down-regulated across all prophase I substages (*p* < 0.05, false discovery rate (FDR), Local test, DESeq2; **Fig 3A**)^33^. Differentially transcribed genes in pachynema relative to leptonema/zygonema were enriched for gene ontologies related to fertilization, positive regulation of transcription by Pol II, and regulation of actin filament polymerization (log_2_fold change > 5.40, DESeq2; **Fig. 3A**). Examples of the 3477 genes down-regulated in diplonema include *Ubl4b* (required for mitochondrial function^34^), *H1fnt* (essential for silencing in spermatids^35^), and *Tssk6* (involved in γH2AX formation^36^) (log_2_ fold change < −2.83, DESeq2; **Fig. 3A**). Pachytene-specific lncRNAs had reported roles in phospholipid translocation, nucleoside diphosphate phosphorylation, and alternative splicing (for example: Gm11837, 1700108J01Rik, and Malat1; **Fig. 3A**)^37^. Thus, we conclude that gene expression profiles for leptonema/zygonema, pachynema, and diplonema have notably different transcription programs.

**Figure 3.**
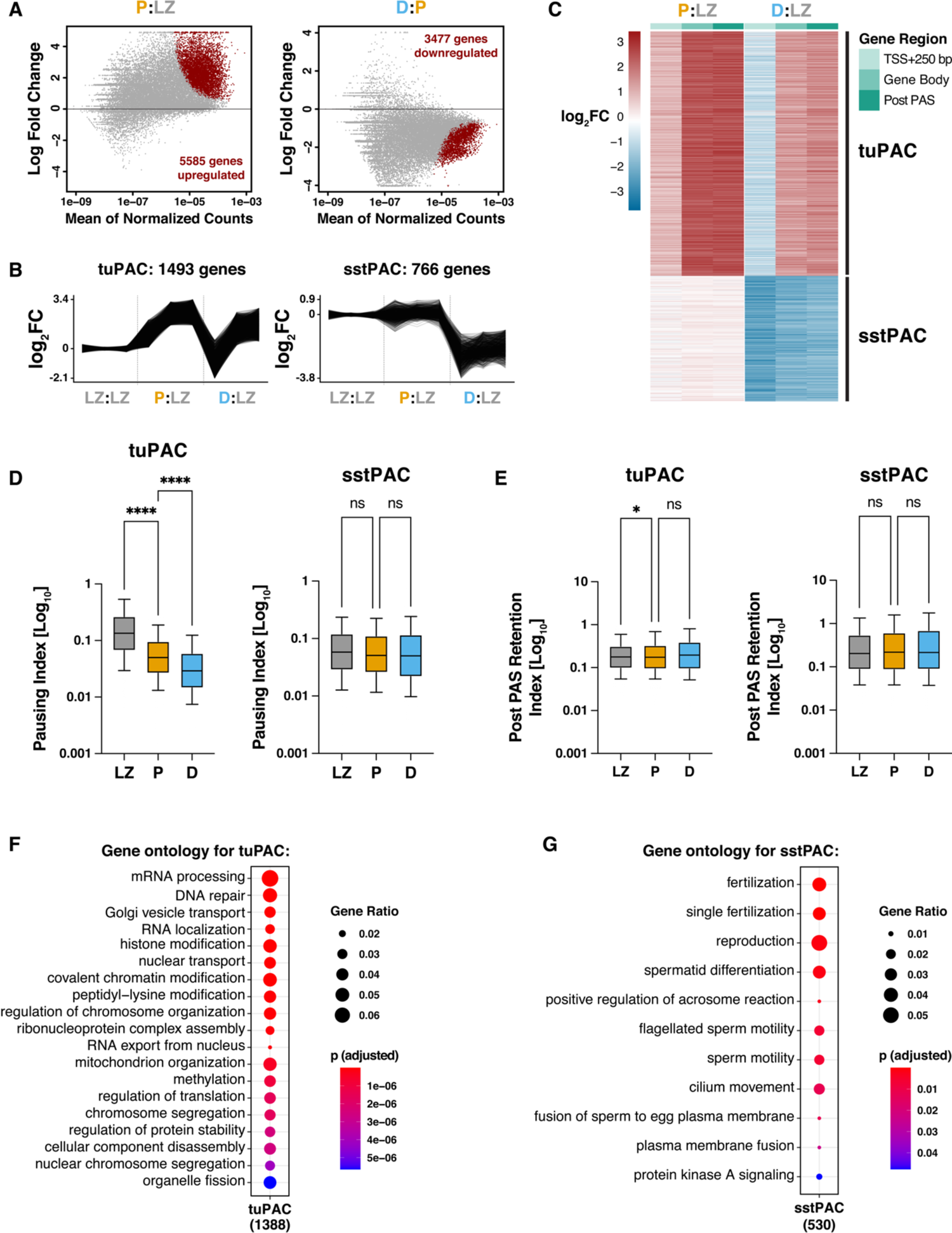
Transcriptional changes between prophase I substages. A) MA plots showing the DEseq2-based differential expression analysis of leChRO-seq read counts within the gene bodies of all annotated genes comparing leptonema/zygonema and pachynema (*left*) and pachynema and diplonema (*right*). B) Trajectories of leChRO-seq read density for individual differentially expressed genes for tuPAC (*top*) and sstPAC (*bottom*) identified in (C). C) Heatmap of log2-transformed fold changes of nascent RNA transcripts enriched or depleted in the selected gene regions for all differentially expressed genes in prophase I identified with DESeq2. Genes were ranked according to changes in leChRO-seq read densities at the TSS + 250 bp, gene body, and post PAS between prophase I substages. Read counts for pachynema (P) and diplonema (D) were normalized to the read counts for leptonema/zygonema (LZ). D) Box and whisker plots of pause index values calculated from the ratio of pause peak density to gene body density of leChRO-seq reads for clusters tuPAC and sstPAC. \*\*\*\**p* < 0.0001, Wilcoxon matched-pairs signed rank test. E) Box and whisker plots of post PAS retention index calculated from the ratio of the leChRO-seq density in the gene body to that in the 3’ end of the gene to 5000 bp downstream for clusters tuPAC and sstPAC. \**p*-value < 0.01, Wilcoxon matched-pairs signed rank test. F-G) Gene ontology analysis for tuPAC (F) and sstPAC (G) from the heatmap in (C). Gene ontology processes are ranked by Benjamini-Hochberg adjusted *p*-value and the ratio of the number of genes in each gene ontology to the number of total genes in each cluster.

To identify distinct patterns of gene expression by the changes in Pol II density observed across prophase I, we clustered DE genes based on their transcription profiles in leptonema/zygonema, pachynema and diplonema according to reads falling within the TSS ± 250 bp (to capture paused Pol II), gene body, and after the polyadenylation cleavage site (PAS) (**Fig. 3B-C**). We identified two major clusters of genes exhibiting differences in Pol II density: *tuPAC (transcriptional <u>u</u>pregulation in <u>pac</u>hynema)* (n = 1493 genes) consisted of genes with a large increase in pachynema, which was partially, but not completely, lost during the transition between pachynema and diplonema, and *sstPAC (<u>s</u>teady-<u>s</u>tate transcription in <u>pac</u>hynema)* (n = 766 genes) which included genes that were largely unchanged in pachynema, but had decreased signal in diplonema (**Fig. 3B-C**). We found that changes at the pause site through prophase I generally increased by a lower amount than the gene body for tuPAC, suggesting that the genes in this cluster are the main targets of genome-wide pause release (**Fig. 3B-C**). Compared to the tuPAC cluster, the genes in the sstPAC cluster did not display promoter-proximal accumulation of Pol II in leptonema/zygonema nor transcriptional bursting in pachynema. To investigate this further, we calculated the pausing index for all genes in clusters tuPAC and sstPAC for leptonema/zygonema, pachynema, and diplonema (**Fig. 3D-E**). Genes in tuPAC exhibited a significantly greater Pol II pausing index in leptonema/zygonema than the genes in sstPAC (**Fig. 3D**; \*\*\*\**p*-value < 0.00001). Moreover, pausing indices at genes in tuPAC decreased during prophase I, whereas those in sstPAC did not change. Taken together, these results suggest genes in sstPAC are transcribed in leptonema/zygonema and pachynema and regulated in a manner that is pause-independent. Conversely, tuPAC genes transcribed in pachynema are activated by the release of Pol II from a paused state that is established in leptonema/zygonema.

To determine the biological function of genes in clusters tuPAC and sstPAC, we performed a gene ontology (GO) analysis (**Fig. 3F-G**). tuPAC genes were enriched in biological process categories with strong relevance for transcriptional regulation (for example: mRNA processing, RNA localization, histone modification, and methylation), post-translational modification (for example: regulation of translation and regulation of protein stability), and meiosis (for example: DNA repair, chromosome segregation, and organelle fission) (**Fig. 3F**). We also found transcripts involved in post-meiotic processes in tuPAC, such as *Ccdc39* (involved in the motility of cilia and flagella), *Spata1* (spermatogenesis associated protein 1), and *Ybx1* and *Brdt* (testis-specific regulators of transcription). Genes in the sstPAC cluster were enriched for post-meiotic GO terms such as spermatid differentiation, sperm motility, and single fertilization (**Fig. 3G**). These results suggest that many of the genes involved in spermatogenesis are transcribed as early as leptonema and undergo a transcriptional decrease in diplonema. This finding may be consistent with reports that meiotic cells transcribe and store mRNAs required for the transcriptionally inert stages of spermatogenesis^13,15^.

### Transcriptional changes lead to changes in mRNA abundance

We next asked whether the dynamic changes in transcriptional activity across prophase I alter mRNA abundance at each stage. As we do not have a way to normalize mRNA-seq data in absolute terms (i.e., the number of transcripts per cell), we focused on examining the correlation between absolute changes in transcription and relative changes in mRNA abundance across prophase I stages. We noted reasonably strong correlations between changes at the level of transcription and mRNA abundance for DE genes (**Fig. S2A-B**; R = 0.54-0.63, Pearson’s correlation). Likewise, within each of the prophase I substages we observed high correlations between the leChRO-seq signal and mRNA abundance for all transcribed genes (**Fig. S2C-E**; Pearson’s R = 0.78-0.82). These results suggest that much of the variation in mRNA abundance during pachynema is driven by the transcriptional burst which peaks in pachynema.

Next, we asked whether mRNA stability may play a more significant role in regulating mRNA abundance as transcription shuts down between pachynema and diplonema. In diplonema, we note that the slope of the line of best fit between leChRO-seq and RNA-seq, which measures the ratio of nascent RNA transcripts to mature mRNA transcripts, decreased by 33% relative to pachynema (pachynema: Pearson’s R = 0.82 ± 0.006; diplonema: Pearson’s R = 0.80 ± 0.006), and the correlation between assays decreased significantly^38^. These findings may indicate that mRNA stability plays a more prominent role during later stages of prophase I as the transcriptional burst in pachynema begins to subside. To put these observations in the context of a specific genomic location, we visualized reads per million (RPM)-normalized RNA-seq libraries at the *Coa3* (Cytochrome C Oxidase Assembly Factor 3), *Cntd1* (Cyclin N-terminal Domain Containing 1), and *Becn1* (Beclin 1; Testis Secretory Sperm-Binding Protein Li 215e) loci (**Fig. S2F**). RNA-seq expression levels for these genes increased from leptonema/zygonema to diplonema, and this pattern was consistent throughout the mouse genome. Together, these observations suggest that transcription is the major determinant of mRNA abundance in leptonema/zygonema and pachynema, but that mRNA stability may become more important in later prophase I stages as transcription begins to shut down.

**Figure S2.**
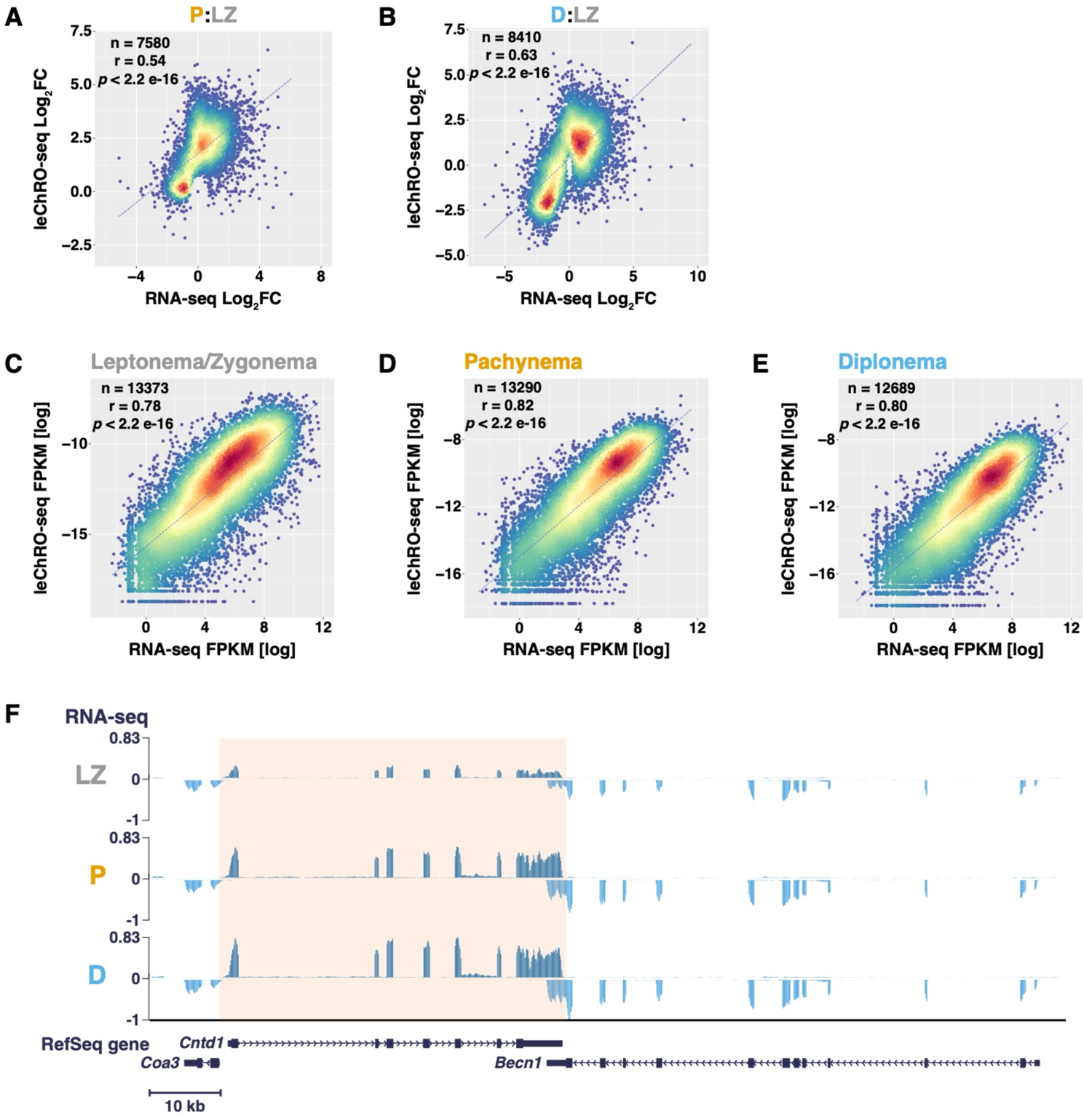
Matched leChRO-seq and RNA-seq libraries are well correlated within individual prophase I substages. A-B) Scatterplots of leChRO-seq vs RNA-seq log_2_FC of all differentially expressed genes for pachynema (A) and diplonema (B) compared to leptonema/zygonema. (See methods section). A-C) Scatterplots of leChRO-seq vs RNA-seq read counts for differentially expressed annotated genes in prophase I cells shown in units of log fragments per kilobase of transcript per million mapped reads (FPKM). (See Methods section). Panels represent scatterplots of matched leChRO-seq and RNA-seq libraries for leptonema/zygonema (A), pachynema (B), and diplonema (C). For each scatterplot, the Pearson correlation coefficient (*r*), number of differentially expressed genes, and *p*-value are shown. Parallel analyses for each prophase I substage revealed a positive correlation between nascent transcription and mRNA expression, with the highest Pearson correlation coefficient in pachynema. D) Reads per kilobase million (RPKM)-normalized RNA-seq signal at the *Cntd1* locus for leptonema/zygonema (LZ; top), pachynema (P; center), and diplonema (D; bottom). mRNA expression levels are highest in diplonema.

### A-MYB coordinates the pachytene transcriptional burst

We asked which transcription factors coordinate the burst of transcriptional activation in pachynema. We identified the location of promoters and enhancers, collectively called transcriptional regulatory elements (TREs), during prophase I by using dREG to identify eRNAs in leChRO-seq data from each prophase I stage^28^. We identified 134,048 TREs during prophase I, the majority of which (65%) were proximal (< 1000 bp) to annotated TSSs, representing candidate promoters (**Fig. S3A**). To identify transcription factors responsible for transcriptional activation in pachynema, we next selected the top TREs that were differentially expressed in leptonema/zygonema compared to pachynema and searched for enriched transcription factor binding motifs (HOMER; **Fig. S3B-E**)^39^. We identified 30 unique motifs with a significant FDR threshold (*p*-value < 1e-54; Fisher’s Exact Test), which we clustered into 5 distinct groups based on similarities in their DNA-binding specificity (**Fig. S3D-E**).

The top scoring motif bound transcription factors (TFs) in the MYB family, including *Myb, A-myb*, and *B-myb* (**Fig. S3E; 4A**), consistent with reports that A-MYB serves as a master regulator in pachynema^19–21,40^. A-MYB ChIP-seq data from pachytene spermatocytes showed a notable overlap (28%) with regulatory elements having MYB binding motifs (Chi-squared test with Yates’ correction, \*\*\**p*-value < 0.0001)^20^. Collectively, our unbiased approach recovered A-MYB as a master regulator of transcriptional changes in pachynema.

Next, we asked whether the genes that exhibit a transcriptional burst during pachynema (tuPAC cluster; see **Fig. 3B-C**) were targets of the A-MYB transcription factor. A-MYB ChIP-seq peaks were strongly enriched in genes in the tuPAC cluster (**Fig. 4B**) — 90% of genes in the tuPAC cluster had A-MYB binding at their promoter, representing a significant enrichment relative to all active genes (\*\*\*\**p* < 0.0001). Genes in the sstPAC cluster were also significantly enriched for A-MYB binding compared with all genes, although the enrichment was significantly less than observed for tuPAC genes. Furthermore, A-MYB target genes were enriched for biological functions reminiscent of tuPAC genes: mRNA metabolic processing, RNA splicing, organelle localization, and histone modification (**Fig. 4C**). Thus, A-MYB is a top candidate to coordinate transcriptional activation in pachynema.

**Figure 4.**
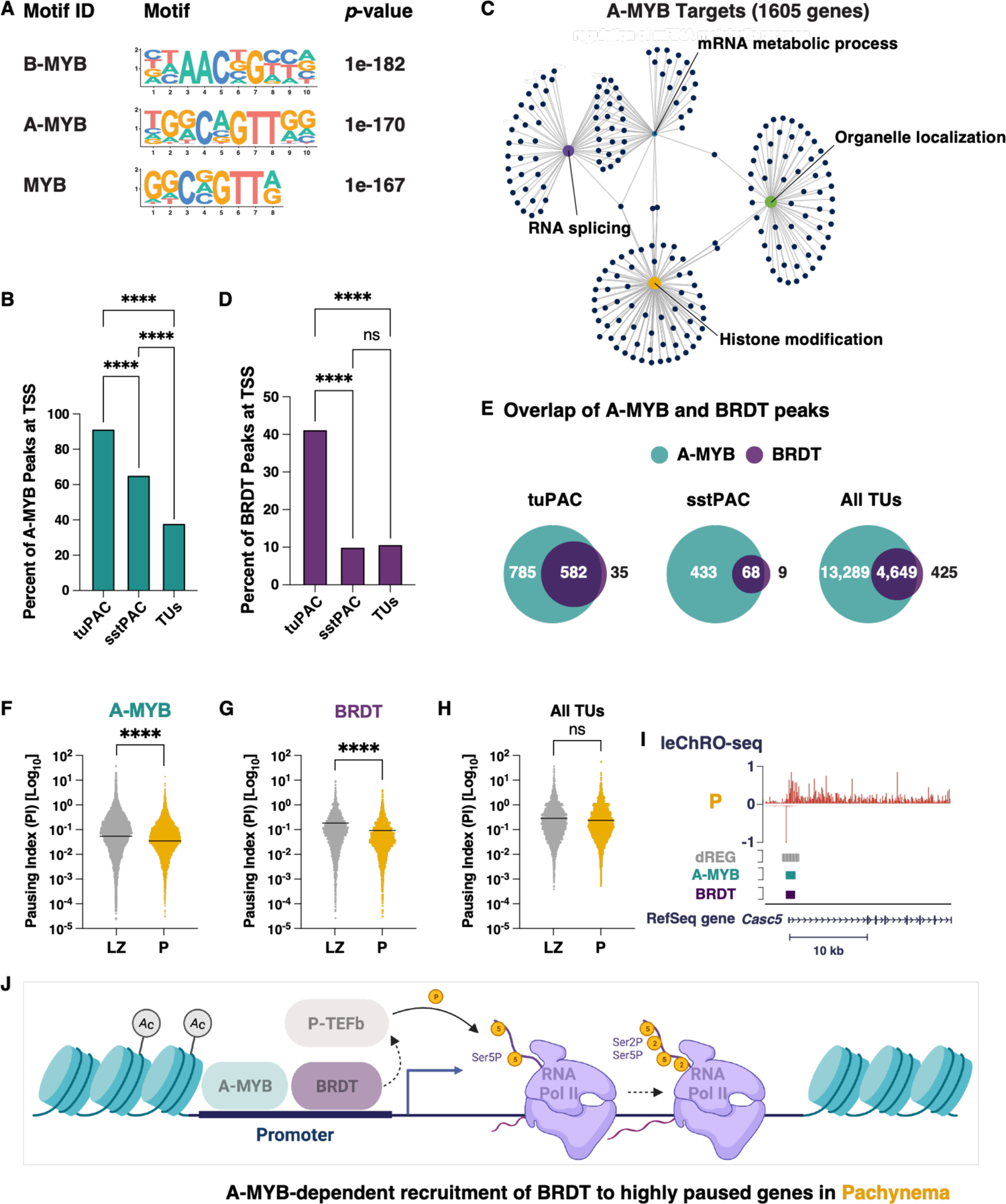
A-MYB and BRDT co-occupy the transcription start sites of differentially expressed genes with higher levels of 5’ Pol II density. A) Transcription factor binding motif enrichment revealed that the MYB transcription factor family displayed the most significant enrichment in differentially transcribed transcriptional regulatory elements in pachytene spermatocytes (FDR < 0.01; Fisher’s Exact Test). The MYB transcription factor binding motifs were identified by HOMER. B,D) The percent of A-MYB (B) or BRDT (D) ChIP-seq peaks overlapping with the transcription start site of genes in tuPAC, sstPAC, or all transcription units. Statistical analyses comparing the percent of A-MYB or BRDT peaks overlapping with the TSSs of genes in tuPAC and sstPAC used the Fisher’s exact test. Statistical analyses comparing the percent of A-MYB or BRDT peaks overlapping with the TSSs of genes in tuPAC or sstPAC and all TUs used the Chi-squared test with Yates’ correction. \*\*\*\**p*-value < 0.00001. C) cnetplot representing the clustering of A-MYB target genes based on gene ontology enrichment analysis. Representative overlap of A-MYB or BRDT ChIP-seq peaks at genes in tuPAC, sstPAC, or all transcription units. F-H) Violin plots of the pausing indices for genes bound by A-MYB or BRDT or not bound by A-MYB or BRDT (All TUs) calculated from the ratio of pause peak density to gene body density of leChRO-seq reads. \*\*\*\**p* < 0.0001, Wilcoxon matched-pairs signed rank test. J) Representative example of A-MYB and BRDT co-binding at the transcription start site of *Casc5*, a gene found in tuPAC. K) A model for the A-MYB dependent recruitment of BRDT to a gene with high levels of 5’ Pol II density in pachynema. We posit that BRDT plays a key role in the regulated release of paused Pol II during prophase I.

**Figure S3.**
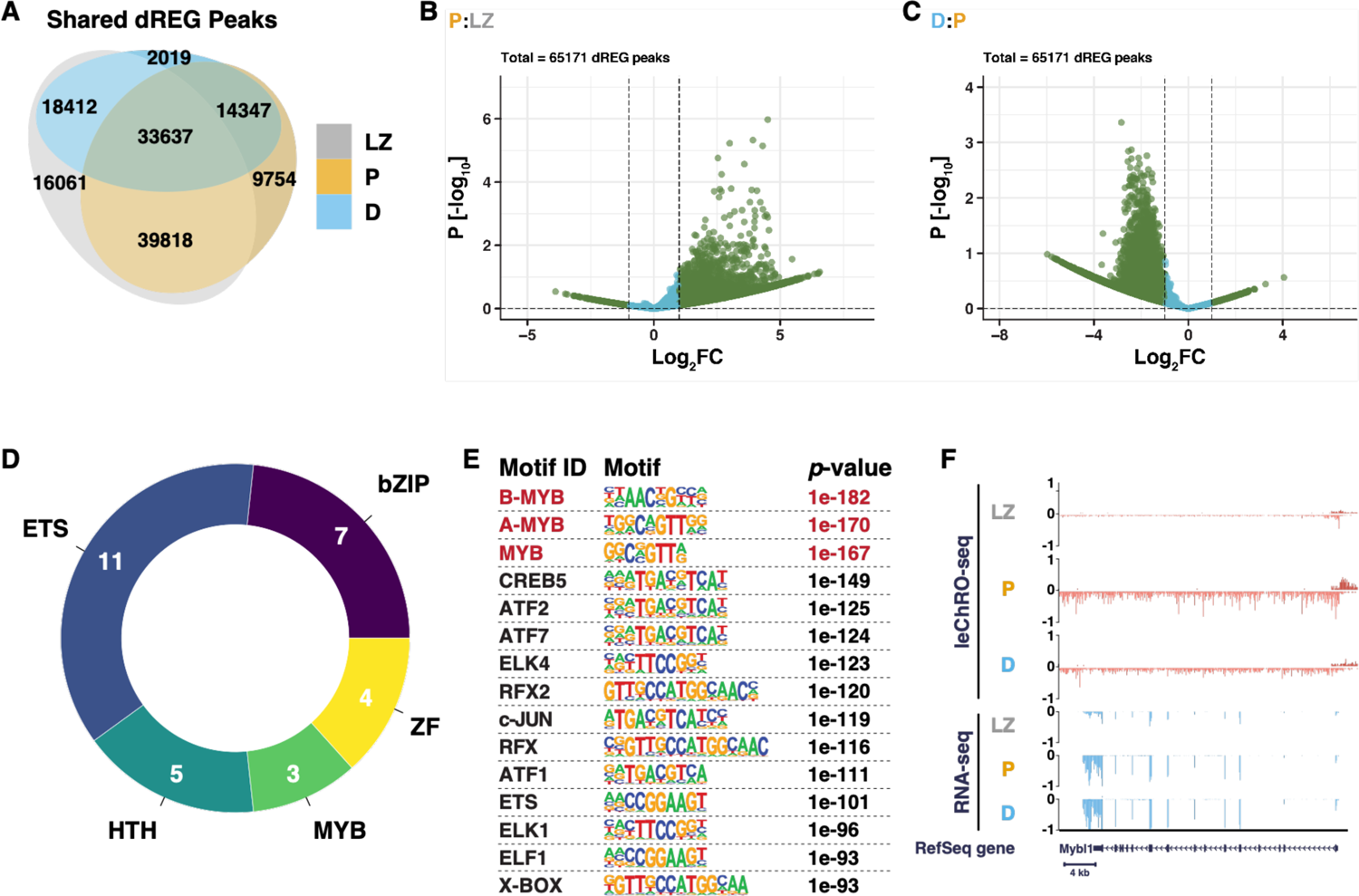
Transcription factors influencing transcriptional activation in pachynema. A) Representative overlap of transcriptional regulatory elements (TREs) identified from dREG by stage. Stage-specific and shared regulatory elements are indicated for each stage of prophase I. Prophase I TREs show considerable overlap between all stages. The number of active TREs decreases throughout prophase I. B-C) Violin plots representing DEseq2-based differential expression analysis of dREG-identified TREs comparing leptonema/zygonema and pachynema (B) and pachynema and diplonema (C). Transcription factor binding motif analysis was performed on the TREs meeting our statistical threshold and are shown in green. D) Donut plot showing the clustering of transcription factor binding motifs based on DNA-binding specificities and transcription factor family. The number of TFs from panel E in each TF family are labeled. E) Transcription factor binding motifs showing the most significant enrichment in upregulated TREs in pachytene spermatocytes. All motifs shown were identified by HOMER and were significantly enriched with an FDR < 0.01 (Fisher’s Exact Test). F) C) leChRO-seq signal and RPKM-normalized RNA-seq signal at the *Mybl1* locus for LZ (top), P (center), and D (bottom).

### A-MYB binding is associated with BRDT binding and pause release

We next sought to determine the mechanism connecting A-MYB with the burst of transcriptional activation in pachynema. Under one model of TF-dependent pause release, a sequence-specific TF recruits transcriptional co-activators that add acetylated histone marks, which recruits the double bromodomain and extra-terminal (BET) family protein, BRD4. BRD4 connects acetylated nucleosomes with the pause-release complex, P-TEFb, to release paused Pol II into productive elongation^43^. Although existing studies have focused on somatic cells, we hypothesized a complementary mechanism exists in germ cells: namely that A-MYB acts to release paused Pol II by recruiting the testis-specific BET protein, BRDT. Therefore, we asked whether A-MYB binding is associated with the recruitment of BRDT at the tuPAC cluster of genes or the sstPAC cluster of gene promoters. We obtained BRDT ChIP-seq data from pachytene spermatocytes^44^, and found that BRDT ChIP-seq peaks were 4.1-fold enriched at TSSs of tuPAC genes compared to either sstPAC or all genes (**Fig. 4D**; \*\*\*\**p*-value < 0.00001). Moreover, we found that >90% of BRDT peaks coincide with A-MYB peaks, regardless of cluster, consistent with the idea that A-MYB works in part by recruiting BRDT (**Fig. 4E**).

To investigate the role of BRDT in transcriptional activation in pachynema, we asked whether BRDT is localized to genes that are paused before high BRDT protein expression in pachynema. The Pol II pausing index in leptonema/zygonema was significantly greater than pachynema for genes bound by A-MYB or BRDT (**Fig. 4F-G**; \*\*\*\**p*-value < 0.00001), consistent with a model in which A-MYB and BRDT released pre-established paused Pol II upon binding in pachynema. In fact, genes that were not bound by either A-MYB or BRDT had no change in pausing index between leptonema/zygonema and pachynema (**Fig. 4H**), demonstrating that A-MYB and BRDT binding sites explain all of the previously observed changes in pausing index genome-wide. These results suggest a model in which A-MYB and BRDT bind to genes having pre-established paused Pol II and facilitate pause release in pachynema (**Fig. 4I-J**).

### A-MYB binding sites poised prior to A-MYB expression in pachynema

We asked whether A-MYB alters chromatin accessibility as its protein binding increases during entry into pachynema. A-MYB protein is reported to reach peak abundance in pachynema^19^, consistent with observations of both transcription and mRNA of the A-MYB-encoding gene, *Mybl1* (**Fig. S3F**). At chromatin accessible regions, ATAC-seq signal peaked in pachynema similarly to nascent transcription (**Fig. 5A-D**). The majority of A-MYB binding sites, however, show a strong ATAC-seq signal in leptonema/zygonema, prior to peak levels of A-MYB protein expression, and retained consistently high levels of accessible chromatin throughout prophase I (**Fig. 5A&E**). Thus, chromatin accessibility at A-MYB binding sites appears to be established prior to peak expression of the A-MYB protein, perhaps through the actions of pioneer transcription factors either in leptonema/zygonema or prior to entry into prophase I. Two candidate pioneer factors include CREB and RFX2, both of which were expressed during spermatogenesis, are essential for entry into meiosis and post-meiotic germ cell differentiation^41,42^, and had binding motifs enriched in transcriptionally active regions in pachynema (**Fig. S3E**, Chi-squared test with Yates’ correction, \*\*\**p*-value < 0.0001). Collectively, our results highlight A-MYB as a transcription factor which binds to poised open-chromatin regions and activates transcription by recruiting BRDT and other pause-release-associated transcriptional machinery.

**Figure 5.**
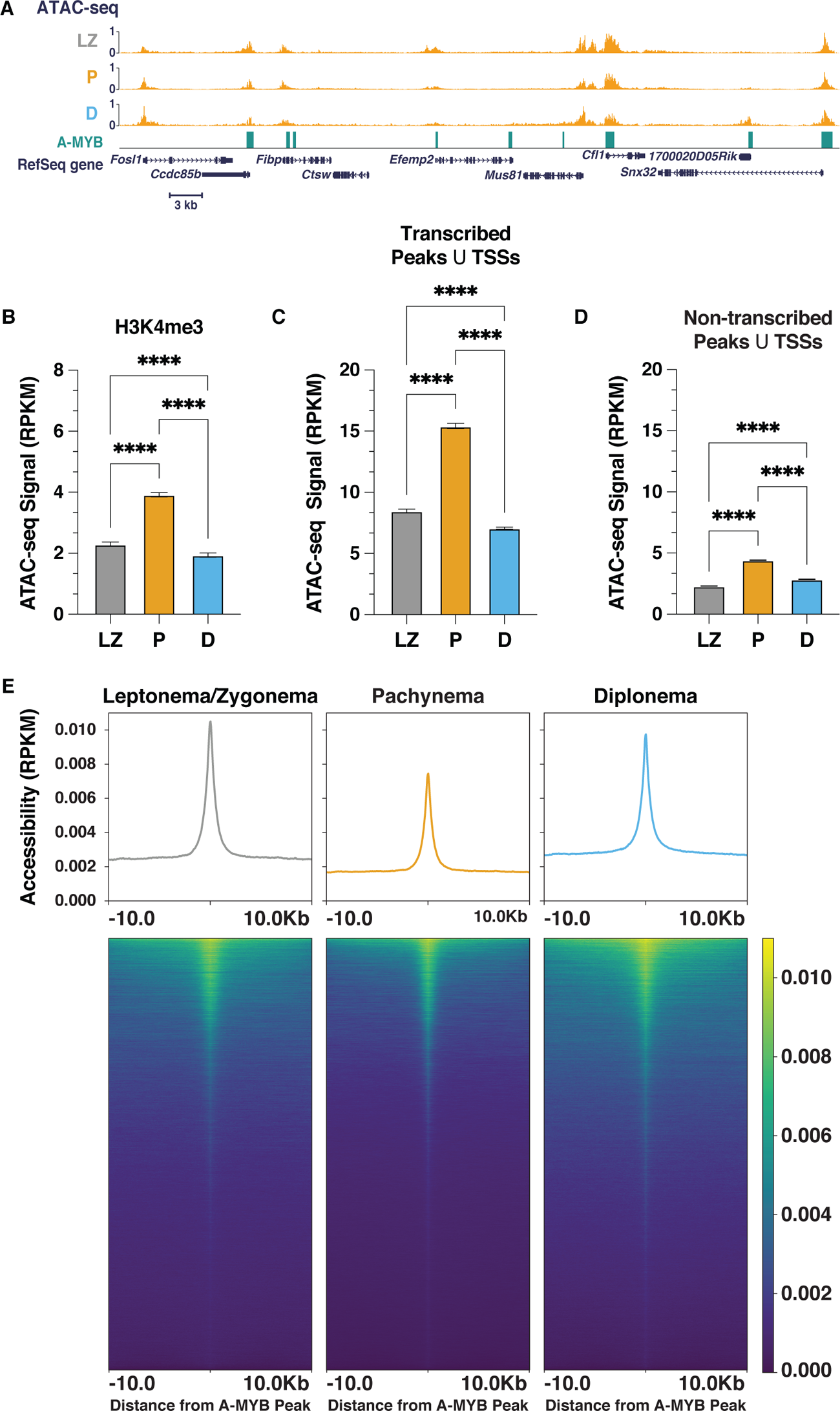
Reprogramming of meiotic chromatin architecture is facilitated by A-MYB binding. A) Reads per kilobase million (RPKM)-normalized ATAC-seq signal at the *Mus81* locus for leptonema/zygonema (top), pachynema (center), and diplonema (bottom). A-MYB ChIP-seq peaks are shown. B-D) Stage-resolved RPKM-normalized ATAC-seq signal at H3K4me3 ChIP-seq peaks (B); transcripted ATAC-seq peaks that overlap with annotated transcription start sites (TSSs) (C); non-transcribed ATAC-seq peaks that overlap with TSSs (D). E) Metaplots (top) and heatmaps (bottom) of RPKM-normalized ATAC-seq signal ± 10Kb from the center of A-MYB ChIP-seq peaks for leptonema/zygonema, pachynema, and diplonema. Peaks are sorted in all prophase I substages by decreasing order of ATAC-seq signal intensity in pachynema.

### Meiotic recombination hotspots have chromatin accessibility and H3K4me3, but no transcription

The designation and distribution of DSB and meiotic recombination hotspots is coordinated by PRDM9^45,46^, a histone methyltransferase that catalyzes the formation of H3K4me3 and H3K36me3 near its binding sites^47–49^. PRDM9 creates a permissive chromatin environment associated with euchromatic and transcriptionally active nuclear compartments^50,51^, all of which are also markers associated with active transcription.

We asked whether DSB hotspots also initiate Pol II transcription. We focused on recombination and DSB hotspots defined by sequenced mouse SPO11 oligos, which provide nucleotide-resolution maps of DSB formation (**Fig. 6**)^47^. As expected, DSBs were located in accessible chromatin, with a signal that peaks in leptonema/zygonema and decreases during the later stages of prophase I (**Fig.6A**). Also as expected, DSBs are also marked by H3K4me3 in leptonema/zygonema (**Fig. 6B**), but lost H3K4me3 in pachynema prior to the burst of transcriptional activation observed above (**Fig. S4A**). However, whereas promoters show a strong signal for transcription that peaks in pachynema (**Fig. 6C**), DSBs show no evidence of transcriptional activity at any point during prophase I (**Fig. 6D**).

**Figure 6.**
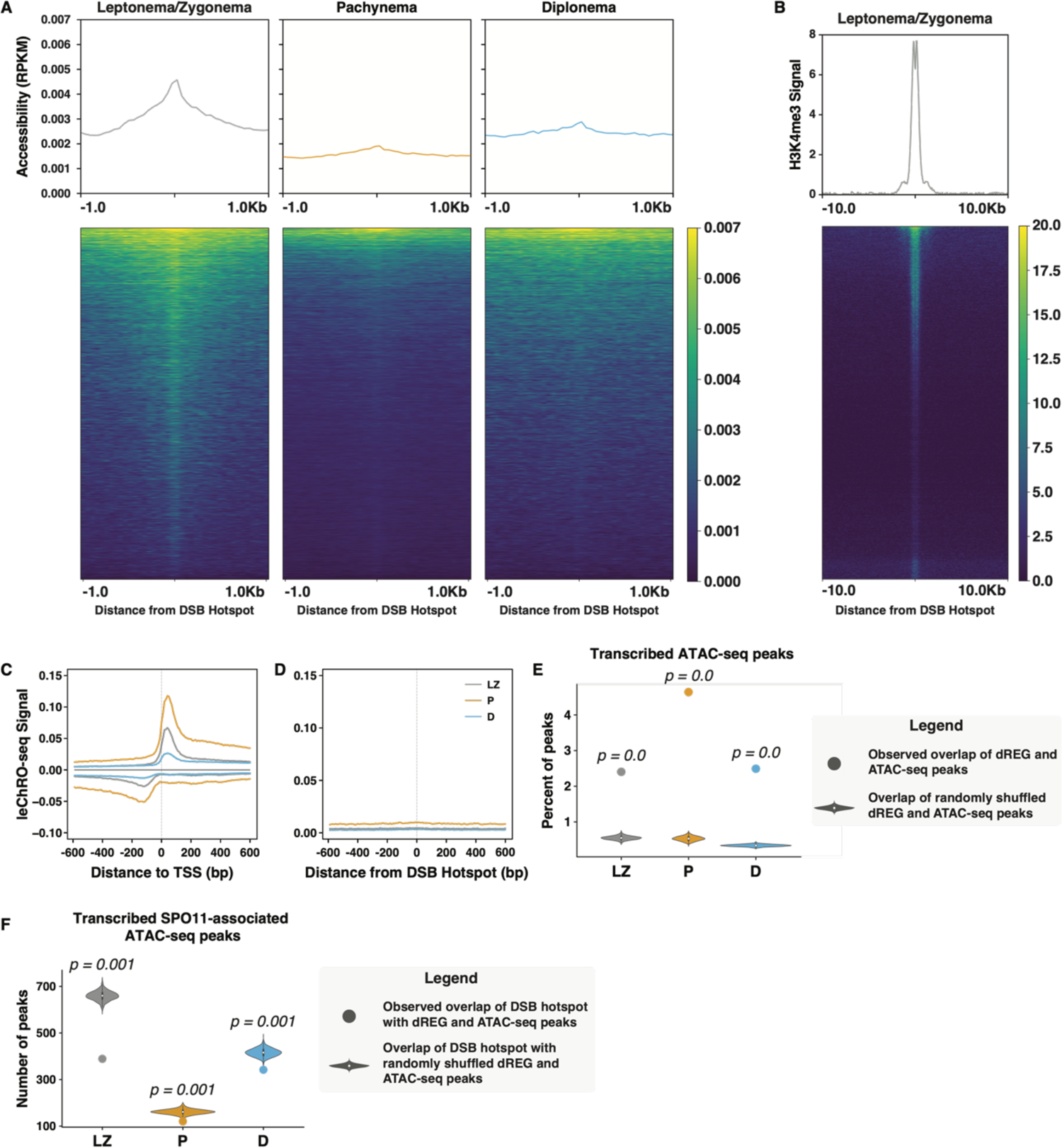
Nascent transcription and chromatin accessibility at DSB and meiotic recombination hotspots. A) Metaplots (top) and heatmaps (bottom) of RPKM-normalized ATAC-seq signal centered on DSB hotspots inferred from SPO11 oligos for leptonema/zygonema (LZ), pachynema (P), and diplonema (D). Peaks are sorted in all prophase I substages by decreasing order of ATAC-seq signal intensity in leptonema/zygonema. B) Metaplot (top) and heatmap (bottom) of H3K4me3 signal centered on DSB hotspots for LZ. C-D) Metaplots of the average leChRO-seq signal centered on annotated TSSs (C) and SPO11 oligos (D) for LZ, P, and D. E) Percentage of observed overlap of dREG and ATAC-seq peaks (*dot*) and overlap of randomly shuffled dREG and ATAC-seq peaks over 1000 iterations (*violin plot*). F) Observed overlap of DSB hotspots inferred from SPO11 oligo data with dREG and ATAC-seq peaks (*dot*) and with randomly shuffled dREG and ATAC-seq peaks (*violin plot*). Empirical *p*-value is reported.

Both DSB formation and active transcription require extensive changes to chromatin that, despite some shared chromatin markers, may not always be compatible. DSBs may therefore be segregated from active promoter and enhancer regions in order to facilitate DSB formation, crossover, and repair. To test this hypothesis, we asked whether transcriptionally active genes and DSBs overlapped more or less than expected by chance after controlling for chromatin accessibility in each prophase-I stage. As expected, ATAC-seq peaks were much more likely to be transcribed, as determined by overlap with dREG, during each stage of prophase I (Empirical *p*-value = 0; **Fig. 6E**). By contrast, DSBs overlapped with active TREs and genes much less than expected by a random model that accounts for chromatin accessibility (**Fig. 6F**; Empirical *p*-value < 0.003). Moreover, we noted no overlap between transcription and two other DSB markers, PRDM9 and DMC1 (**Fig. S5A-B**), supporting a lack of transcription at DSBs regardless of the marker. We conclude that although DSBs and TREs share many chromatin features, they rarely overlap, perhaps so that they are each able to specialize in a distinct molecular function.

**Figure S4.**
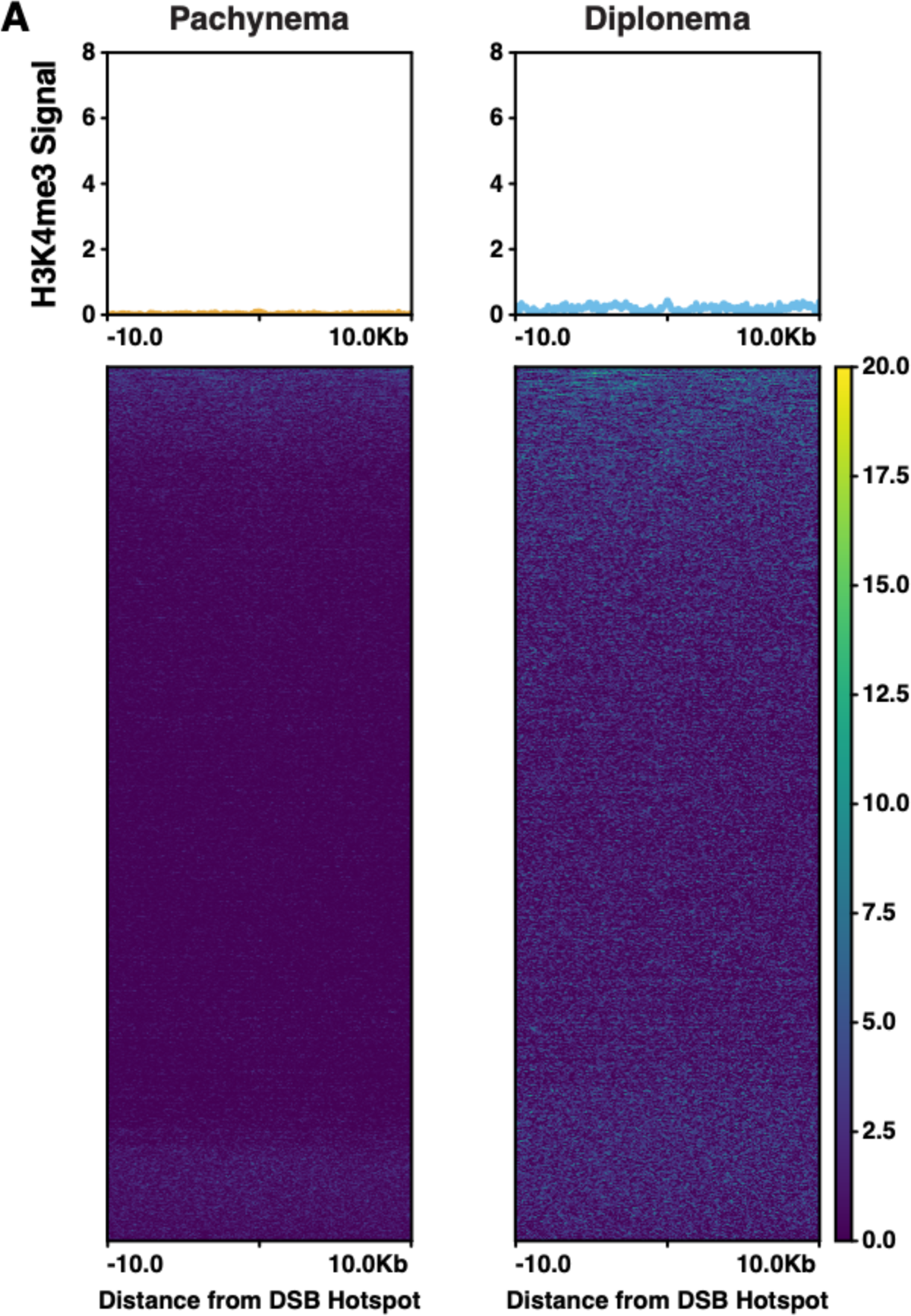
A) Metaplots (top) and heatmaps (bottom) of H3K4me3 signal centered on DSB hotspots, inferred from SPO11 oligo data, for pachynema and diplonema.

**Figure S5.**
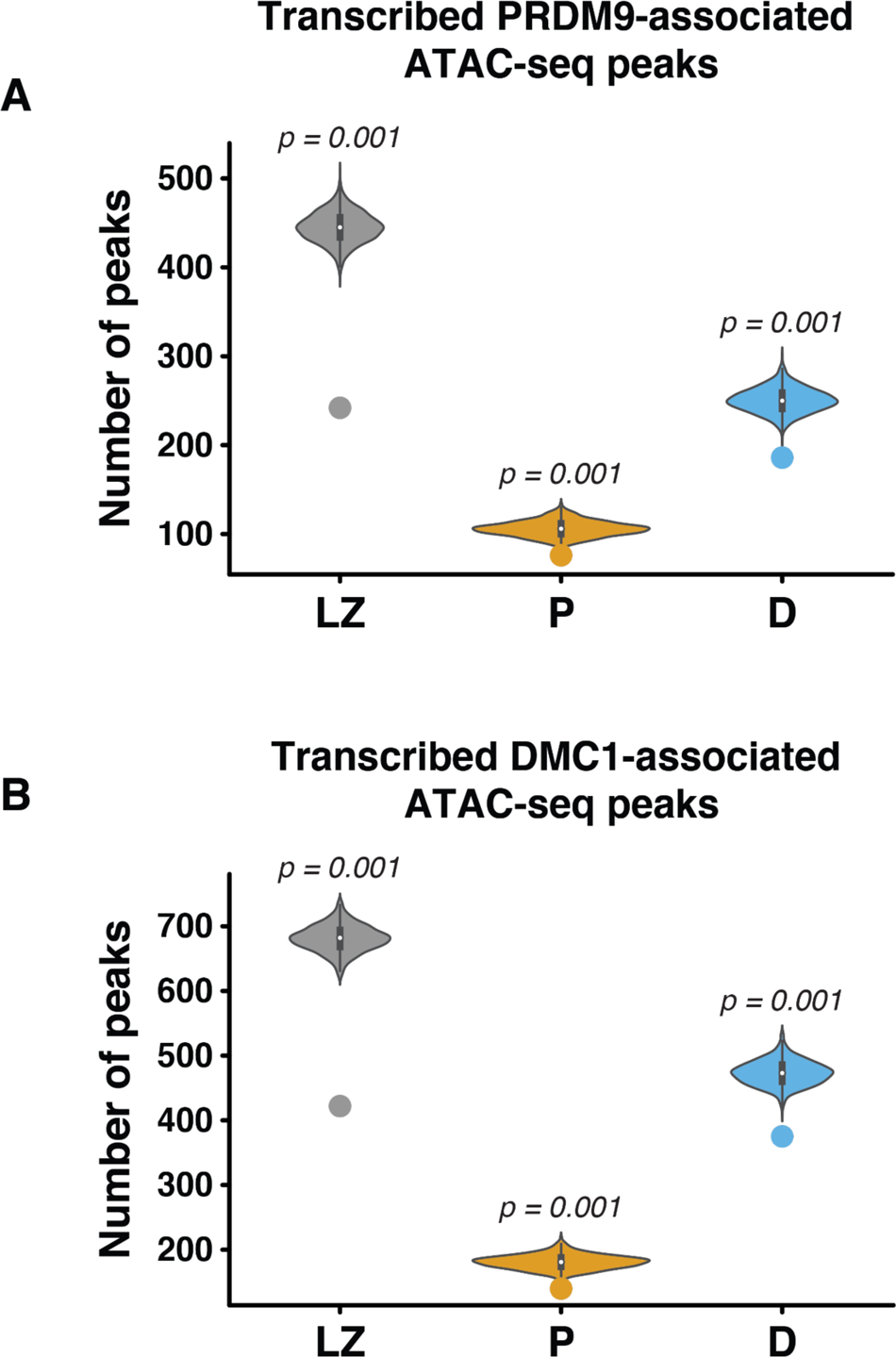
A-B) Observed overlap of PRDM9 motif (A) and DMC1 peaks (B) with dREG and ATAC-seq peaks (*dot*) and with randomly shuffled dREG and ATAC-seq peaks (*violin plot*). Empirical *p*-value is reported.

## DISCUSSION

Prophase I spermatocytes require extensive reprogramming of gene expression to support the transition into meiosis and then to accomplish the morphological transformation of haploid germ cells during spermiogenesis^8,11,13,14,21,50,52–54^. Although meiotic chromosomes are highly organized by the synaptonemal complex, prophase I cells maintain an extremely diverse, complex, and tightly regulated transcriptional program^11,50,52,53^. While recent advances in high-throughput sequencing have identified the overall structure of meiotic chromosomes and transcription hub formation, very little information is available on the coordinated recruitment of the transcription machinery to meiotic chromatin during discrete substages of prophase I. Here, we used leChRO-seq to directly map the presence of transcriptionally competent RNA Polymerase II (Pol II) at single nucleotide resolution for all meiotic prophase I substages from leptonema to diplonema (**Fig. 2**). We found that the transcriptional activity of Pol II is nearly three times greater in pachynema than leptonema/zygonema, diplonema, and round spermatids (**Figs. 1-2**). We provide evidence that Pol II pausing during early elongation in leptonema/zygonema is a general feature of the transcription cycle in prophase I and is connected to the rapid and synchronous activation of transcription of thousands of genes in pachynema (**Fig. 2**). Indeed, the differentially expressed (DE) genes most strongly associated with pausing encode essential regulators of male germ cell differentiation, including those required for meiotic progression, sperm motility, and testis-specific transcription factors (**Fig. 3**). These observations suggest that Pol II pausing provides a mechanism to finely tune meiotic genes to distinct regulatory cues and execute stringent spatiotemporal control of transcription during prophase I.

Our study has shown that A-MYB and BRDT mediate Pol II pause-release at differentially upregulated genes during pachynema. We demonstrated that A-MYB and BRDT localization in pachynema was significantly associated with actively transcribed genes with higher Pol II pausing indices in leptonema/zygonema (**Fig. 4**). Given that A-MYB binding co-occurs with H3K27ac at active regulatory elements during meiosis, we propose that BRDT is recruited to A-MYB-bound promoters through its recognition of acetylated lysine residues. Since the C-terminal domain of BRDT has been shown to interact with components of the P-TEFb complex, BRDT is likely critical for regulating Pol II pause-release and transcriptional activation of A-MYB-bound genes during prophase I. However, we cannot exclude the possibility that other BET proteins or TFs influence the rate of pause-release in prophase I spermatocytes. Resolving the network of TFs and developmental cues responsible for Pol II pausing and transcription elongation control during prophase I provides an exciting future research direction.

One key feature of mammalian prophase I chromosomes is the spatial clustering of highly transcribed loci even as topologically associated domains (TADs), or long-range contacts, are lost in pachynema^11,50,52,53^. Thus, to determine the temporal establishment of transcriptional regulatory elements (TREs) during prophase I, we profiled active TREs at high resolution using a comparative analysis of nascent transcription and chromatin accessibility (**Figs. S3 and 5**). Our data suggest that the majority of prophase I TREs are established upon entry into meiosis. Regulatory element usage in leptonema/zygonema coincides with the global accumulation of promoter-proximal Pol II and, intriguingly, paused Pol II has been shown to have a crucial role in maintaining accessible promoter chromatin architecture prior to and independently from gene activation^55^. In agreement with this observation, we found a nearly two-fold increase in genome-wide chromatin accessibility between leptonema/zygonema and pachynema at all gene TSSs (**Fig. 5**). Given that meiotic cells undergo profound chromosome compaction during the hallmark events of prophase I, these results suggest that chromatin accessibility is not mechanistically coupled with 3D genome reorganization at the pachytene stage. We posit that gene-specific and local chromatin accessibility in pachynema is likely mediated by Pol II pausing, resulting in a permissive environment that enhances transcription by preventing nucleosome assembly at active promoters in pachynema.

The programmed formation of hundreds of DNA double-strand breaks (DSBs) is essential for initiating meiotic recombination and ensuring proper fertility^2^. In mice and humans, the distribution of meiotic DSBs is controlled by the binding of PRDM9 and SPO11^2,56^. To further explore the control of gene expression at DNA DSBs and recombination hotspots, we overlaid our profiles of chromatin accessibility and nascent transcription with previously published maps of PRDM9, SPO11, and DMC1 binding in C57Bl/6 males^47,51^. In particular, we found that DSB and recombination hotspots most frequently occur at regions of accessible chromatin, likely as a result of the histone methyltransferase activity of PRDM9^57^. However, engaged Pol II was undetectable at sites of meiotic recombination in pachynema, which stands in stark contrast to the widespread transcriptional activity of meiotic chromosomes at this stage. Therefore, it is conceivable that a mechanism of active transcriptional repression exists at sites of DNA DSBs and eventual crossovers or that PRDM9, SPO11, and/or DSB repair complexes actively repel the transcriptional machinery that is activated in pachynema.

In summary, we report the first comprehensive profiling of Pol II occupancy genome-wide in prophase I spermatocytes. Our study directly demonstrates the global presence of paused Pol II at the TSSs of highly expressed genes in early prophase I, thus providing a mechanism for the temporal control of transcription during spermatogenesis. We examined active *cis*-regulatory elements and identified multiple TF pathways associated with the massive transcriptional activation observed in pachynema. We show that BRDT is recruited to highly paused genes in pachynema and is an important transcriptional co-activator of A-MYB-bound genes. Our observations of dynamic gene expression during prophase I provide new insights into transcriptional regulation during key meiotic events, such as DSB initiation and repair. Indeed, these data represent an essential resource for the discovery of previously unknown regulatory elements and TFs coordinating the developmental transitions of spermatogenesis and ensuring successful male fertility.

## MATERIALS AND METHODS

### Care and use of experimental animals

The experiments described herein used mice on the C57Bl/6J background, obtained from Jackson Laboratories. All mice were housed under strictly controlled conditions of temperature and light:day cycles, with food and water *ad libitum*. All mouse studies were conducted with prior approval by the Cornell Institutional Animal Care and Use Committee, under protocol 2004-0063.

### Chromosome spreading and immunofluorescent staining

Prophase I chromosome spreads and immunofluorescent staining were prepared as previously described^58,59^. Testis tubules were incubated in hypertonic elution buffer (30mM Tris-HCl pH 7.2, 50 mM sucrose, 17 mM trisodium dihydrate, 5 mM EDTA, 0.5 mM DTT, 0.1 mM PMSF, pH 8.2-8.4) for one hour. Small sections of tubules were minced in 100 mM sucrose solution and spread onto 1% Paraformaldehyde, 0.15% Triton X-100 coated slides. Slides were incubated in a humid chamber for 2.5 hours at room temperature. Slides were dried for 30 minutes, washed in 0.4% Photoflo (KODAK, Geneva NY) diluted in PBS (800 ml in 200 mL PBS), 0.1% Triton X-100 diluted in PBS (1 mL and 199 mL PBS) and blocked in 10% antibody dilution buffer (ADB:3% bovine serum albumin, 0.05% Triton in 1 x PBS) diluted in PBS (20 mL and 180 mL). Primary antibodies used included rabbit-anti-SYCP3 (custom-made) ^60^, mouse-anti-RNA Pol II (Millipore Sigma #05-623; 1:2000 dilution), rat-anti-RNA Pol II Ser2P (Millipore Sigma # 04-1571; 1:1000 dilution), rat-anti-RNA Pol II Ser5P (Millipore Sigma #04-1572; 1:500 dilution), and anti-A-MYB (Sigma Prestige Antibodies #HPA-008; 1:1000 dilution). Primary antibodies were diluted in antibody dilution buffer, spread across the surface of the slide, and incubated at 4°C overnight. Slides were washed for 10 minutes each in 0.4% Photoflo, 0.1% Triton X, and 10% antibody dilution buffer. Alexafluor(tm) secondary antibodies (Molecular Probes Eugene OR, USA) were used for immunofluorescent staining at 37°C for one hour. Secondary antibodies were diluted in antibody dilution buffer and spread in a similar fashion to the primary antibodies. Slides were washed as previously described, dried, and mounted with Prolong Gold antifade (Molecular Probes). All secondary antibodies were raised specifically against F_c_fraction, F_ab_-fraction purified and conjugated to Alexafluor 488 or 594.

### Image acquisition

Images were acquired using a Zeiss Axiophot Z1 microscope attached to a cooled charge-coupled device (CCD) Black and White Camera (Zeiss McM). The images were captured and pseudo-colored by means of ZEN 2 software (Carl Zeiss AG, Oberkochen. Germany). Exposure time was consistent between antibodies, cells, and mice. Brightness and contrast of images were adjusted using ImageJ (National Institutes of Health, USA) after fluorescence intensity measurements were calculated.

### FIJI ImageJ macro script for RNA Pol II, RNA Pol II Ser2P, and RNA Pol II Ser5P fluorescence intensity measurements

A FIJI macro script was created using the available tools in FIJI. Images were in .czi file format, with DAPI in blue; SYCP3 in green; and RNA Pol II, RNA Pol II Ser2P, and RNA Pol II Ser5P in red. Intensities of DAPI, RNA Pol II, RNA Pol II Ser2P, and RNA Pol II Ser5P were determined using Otsu’s thresholding; intensities of RNA Pol II, RNA Pol II Ser2P, and RNA Pol II Ser5P were normalized to DAPI signal intensity to account for the DNA content of each cell. The script used was as follows:

~~~
dir=getDir(“Choose the directory containing czi files”);
outfile=substring(dir,0,dir.length-1);
outfile=outfile+”.txt”;
files=getFileList(dir);
run(“Set Measurements…”, “mean limit redirect=None
decimal=2”);
//Array of averages for each channel
Dave=newArray(3);
fp=File.open(outfile);
for (k = 0; k < files.length; k++) {
            run(“Bio-Formats”, “open=[“+dir+files[k]+”]
color_mode=Grayscale quiet view=Hyperstack stack_order=XYCZT”);
            for (i = 0; i < 3; i++) {
                        Stack.setPosition(i+1,1,1);
                        setAutoThreshold(“Otsu dark”);
                        run(“Measure”);
                        Dave[i]=getResult(“Mean”);
                        }
print(fp,files[k]+”\t”+Dave[0]+”\t”+Dave[1]+”\t”+Dave[2]);
            close();
            }
File.close(fp);
~~~

### Quantification and statistical analysis of images

Statistical analyses of fluorescent intensity displayed in **Fig. 1** were completed using GraphPad Prism v9.0 for Macintosh (GraphPad Software, San Diego California USA, www.graphpad.com). Mean values ± 95% confidence interval (CI) were presented and alpha values were established at 0.05. All statistical analyses performed utilized One-way ANOVA with the post-hoc Tukey’s multiple comparison’s test. Images of diakinesis-staged cells were not included in the statistical analysis due to low sample size.

### Isolation of mouse spermatogenic cells by STAPUT

Testes from adult wildtype mice (day 70-80 pp; n = 40) were extracted and decapsulated prior to enrichment of prophase I staged cell types using the gravitational cell separation method, STAPUT, which is based on separation by cell diameter and density at unit gravity^59,61^. Purity of collected fractions was determined by staining against proteins defining prophase I substages, specifically SYCP3 and γH2AX (EMD Millipore 05-636), and imaged on a Zeiss Axiophot with Zen 2.0 software. Leptotene and zygotene, pachytene, and diplotene cells were at approximately 87%, 84%, and 81% purity, respectively, with potential contamination from spermatogenic cells of either earlier or later developmental timing. Round and elongating spermatids were identified by DAPI staining and collected at l00% enrichment. Cells isolated by STAPUT were used as input for leChRO-seq (n = 4) and RNA-seq (n = 3) library preparation.

### Isolation of mouse spermatogenic cells by Flow Cytometric Analysis and Fluorescence Activated Cell Sorting (FACS)

Testes from adult wildtype mice (day 70-80 pp, n = 2) were extracted and decapsulated prior to enrichment of prophase I staged cell types using FACS ^62^. Prior to sorting, testicular single cells were stained with Hoechst dye and Propidium Iodide for selection based on DNA content and dead cell exclusion, respectively. Data analysis was done using the BD FACSDiva Software v6.1.3 on a FACSAria II cell sorter. Hoescht was excited using the 355 nm laser. Sorting flow rate was adjusted to 7-32 events/second. Purity of collected cell populations was accessed as described previously for STAPUT. Leptotene and zygotene, pachytene, and diplotene cells were at approximately 80%, 71%, and 93% enrichment, respectively. Nuclei from cells isolated by FACS were used as input for ATAC-seq libraries (n = 2).

### Chromatin isolation for nuclear run-on assays

The methods of chromatin isolation were based on work described by Chu et al., 2018^23^. For chromatin isolation from STAPUT-sorted cells, we added 1 ml of 1X NUN buffer (0.3 M NaCl, 1M Urea, 1% NP-40, 20 mM HEPES, pH 7.5, 7.5 mM MgCl_2_, 0.2 mM EDTA, 1mM DTT, 50 units per ml RNase Cocktail Enzyme Mix (Ambion, AM2286), 1x Protease Inhibitor Cocktail (Pierce, A32965). Samples were vigorously vortexed for 1 min. 500 µl of NUN buffer was added to each sample and vortexed for another 30 s. The samples were then incubated in a Thermomixer C at 12°C, 1,500 rpm, for 30 min. Chromatin samples were centrifuged at 12,500g for 30 min at 4°C. Following removal of the NUN buffer from the chromatin pellet, the chromatin pellet was washed with 1 ml 50 mM Tris-HCl, pH 7.5, with 40 units per ml RNase inhibitor, centrifuged at 10,000g for 5 min at 4°C, at which point the buffer was discarded for a total of three washes. After washing, 50 ul of chromatin storage buffer (50 mM Tris-HCl, pH 8.0, 25% glycerol, 5 mM Mg(CH_3_COO)_2_), mM EDTA, 5 mM DTT, 40 units per ml RNase inhibitor) was added to each sample. The samples were sonicated with a Bioruptor on high power, with a cycle time of 10 min with cycle durations of 30 s on and 30 s off. The sonication was repeated up to four times as needed to get the chromatin pellet into suspension. The samples were stored at −80°C.

### Nuclear run-on assay using radioactive [α32P]CTP

Chromatin was isolated from STAPUT-sorted fractions containing either mixed populations of prophase I cells (n = 17) or round and elongated spermatids (n = 4). Prior to the nuclear run-on assay, DNA content of isolated chromatin was measured to ensure equal input of material per sample (500 ng of DNA). Three 10-fold serial dilutions of chromatin isolated from two Jurkat samples was also used to standardize each nuclear run-on experiment to account for radioactive decay of [α32P]CTP. The nuclear run-on was started by mixing isolated chromatin with 50 ml 2x chromatin run-on buffer (10 mM Tris-HCl pH 8.0, 5 mM MgCl_2_,1 mM DTT, 300 mM KCl, 400 µM ATP (NEB N0450S), 400 µM GTP (NEB N0450S), 400 µM UTP (NEB N0450S), 40 mM 32P-CTP (Perkin Elmer BLU008H), 0.8 units/µl SUPERase in RNase Inhibitor (Life Technologies AM2694), and 1% Sarkosyl (Fischer Scientific #AC612075000)). The run-on reaction was incubated at 37°C for 10 minutes and then stopped by adding Trizol LS (Life Technologies 10296-101). RNA was extracted with 1-bromo-3-chloropropane, pelleted with GlycoBlue treated water, and washed with 100% and 75% ethanol. The RNA was resuspended in diethyl pyrocarbonate (DEPC) H_2_0 and any free nucleotides were captured by the use of two P-30 columns. All samples were assessed for radioactive levels on an LS Beta Counter. The relative transcriptional activity of leptonema/zygonema (LZ), pachynema (P), and diplonema (D) was calculated using the formula below:

1. *Calculate line of best fit for observed CPM of Jurkat standards*: *y = mx + b*, where *y =* rate of ionization events per minute (CPM); and *x =* DNA concentration
2. *Calculate expected CPM for samples using line of best fit:* substitute *x* with measured DNA concentration of sample
3. *Relative transcriptional activity:*

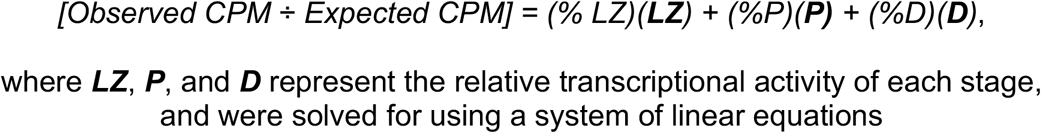

The relative transcriptional activity of round and elongated spermatids, enriched at 100%, was calculated from the *[Observed CPM ÷ Expected CPM]* ratio. Mean values ± standard deviation (SD) of the relative transcriptional activity for each cell type were presented in Fig. 1 and plotted using GraphPad Prism. Statistical analyses performed utilized One-way ANOVA with the post-hoc Tukey’s multiple comparison’s test. Alpha values were established at 0.05.

### leChRO-seq library preparation

leChRO-seq libraries were prepared as previously described by Chu et al., 2018 ^23^. Briefly, chromatin from enriched prophase I substages was mixed with 50 µl 2x run-on master mix (10 mM Tris-HCl pH 8.0, 5 mM MgCl_2_,1 mM DTT, 300 mM KCl, 10 mM Biotin-11-CTP (Perkin Elmer # NEL542001EA), 400 uM ATP (NEB N0450S), 0.8 uM CTP (NEB N0450S), 400 uM GTP (NEB N0450S), 400 uM UTP (NEB N0450S), 40 uM Biotin-11-CTP (Perkin Elmer # NEL542001EA), 40 units SUPERase in RNase Inhibitor (Life Technologies AM2694), and 1% Sarkosyl (Fischer Scientific #AC612075000). The run-on reaction was incubated at 37°C for 5 minutes and then stopped by adding Trizol LS (Life Technologies 10296010). RNA was extracted by adding chloroform and precipitating the aqueous phase with GlycoBlue and ethanol. Unincorporated biotin nucleotides were removed with Micro Bio-Spin P-30 Gel Columns, Tris Buffer (RNase-free) (Bio-Rad #7326250). Nascent RNA was purified by binding streptavidin beads (NEB S1421S) and washed as described by Mahat et al., 2016^63^. The RNA was removed from beads by Trizol and then followed with the 3’ adapter ligation using T4 RNA Ligase 1 (NEB M0204L). After this, a second bead binding was performed followed by a 5’ de-capping using RppH (NEB M0356S). The 5’ end of the transcript was phosphorylated using PNK (NEB M0201L) and then purified with Trizol (Life Technologies 15596-026). After this, a 5’ adapter was ligated onto the RNA transcript using T4 RNA Ligase I (NEB M0204L), and a third bead binding was performed. RNA was purified again with a Trizol and ethanol extraction. A reverse transcription reaction was performed in order to generate cDNA using SuperScript IV Reverse Transcriptase (Life Technologies 18090010). The cDNA was amplified with Q5 High-Fidelity DNA Polymerase (NEB M0491L) in order to generate the leChRO-seq libraries which were prepared using manufacturer’s protocol (Illumina) and paired-end sequenced by means of Illumina NextSeq500 at Cornell University Biotechnology Resource Center.

### Mapping of leChRO-seq sequencing reads

In order to align leChRO-seq data, a publically available script was used (https://github.com/Danko-Lab/utils/tree/master/proseq). The libraries were prepared by means of adapters containing a specific molecular identifier. For these specific transcripts, PCR duplicates were removed using PRINSEQ lite^64^. The adapters were cut from the 3’ end of remaining reads using cutadapt with a 10% error rate^65^. Reads from this data were mapped using BWA^66^ to the mouse reference genome (mm10) in addition to a single copy of the Pol I ribosomal RNA transcription unit (GenBank ID #U13369.1). The specific location of the RNA polymerase active site was described by means of a single base which denoted the 5’ end of the nascent RNA. This information corresponded to the position of the 3’ end of each sequenced read. Mapped reads were converted to bigWig format using BedTools^67^. Libraries were normalized using the following formula:

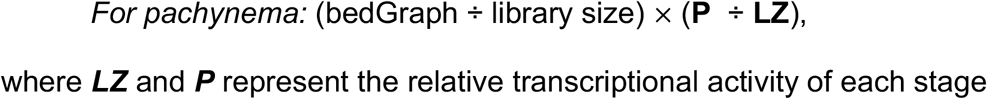

This formula was repeated for all prophase I substages. The WashU Epigenome Browser was used for visualization of leChRO-seq libraries^68^.

### RNA extraction and RNA-seq library preparation

Immediately following STAPUT, testicular cells were lysed with TrizolLS (Thermo Fisher). Total RNA was purified using TrizolLS according to the commercial protocol with the following additions: after the first phase separation, an additional chloroform extraction step of the aqueous layer was performed using Phase-lock Gel heavy tubes (Quanta Biosciences); addition of 1ul Glyco-blue (Thermo Fisher) immediately prior to isopropanol precipitation; two washes of the RNA pellet with 75% ethanol. If the RNA integrity results indicated co-purified genomic DNA, it was removed with the RapidOUT DNA Removal kit (Thermo Fisher). RNA sample quality was confirmed by spectrophotometry (Nanodrop) to determine concentration and chemical purity (A260/230 and A260/280 ratios) and with a Fragment Analyzer (Advanced Analytical) to determine RNA integrity. PolyA+ RNA was isolated with the NEBNext Poly(A) mRNA Magnetic Isolation Module (New England Biolabs). TruSeq-barcoded RNAseq libraries were generated with the NEBNext Ultra II Directional RNA Library Prep Kit (New England Biolabs). Each library was quantified with a Qubit 2.0 (dsDNA HS kit; Thermo Fisher) and the size distribution was determined with a Fragment Analyzer (Advanced Analytical) prior to pooling. Libraries were sequenced on a NextSeq500 instrument (Illumina). At least 20M single-end 75bp reads were generated per library.

### Mapping of RNA-seq sequencing reads

Reads were trimmed for low quality and adaptor sequences with cutadapt v1.8; parameters: -m 50 -q 20 -a AGATCGGAAGAGCACACGTCTGAACTCCAG --match-read-wildcards. Reads were mapped to the reference genome (mm10.rRNA.fa.gz) using STAR v2.5.3a^69^. Mapped reads were converted to bigWig format using BedTools. To normalize for sequencing depth, libraries were scaled using reads per million mapped reads (RPM). The WashU Epigenome Browser was used for visualization of RNA-seq libraries.

### ATAC-seq library preparation

This methodology is directly based on the original design^70, 71^.

1. *Prepare nuclei:* In order to prepare nuclei, 50,000 cells isolated by FACS were spun at 500g for 5 minutes immediately after sorting, followed by a wash using 50µl of cold 1 x PBS and centrifugation at 500g for 5 minutes. Cells were then lysed using a cold lysis buffer (10mM Tris-HCl, pH 7.4, 10mM NaCl, 3mM MgCl_2_ and 0.1% IPEGAL CA-630). Immediately following lysis, nuclei were spun at 500g for 10 minutes in a refrigerated centrifuge. In order to avoid cell loss during nuclei preparation, a fixed-angle centrifuge was used and cells were carefully pipetted away from the pellet after centrifugation. Nuclei were stored at −80°C.
2. *Transpose and purify*. Stored nuclei were thawed on ice and the pellet was resuspended in a transposase reaction mixture (25µl 2 x TD buffer, 2.5µl transposase (Illumina) and 22.5µl nuclease-free water). The transposition reaction mixture was carried out for 30 minutes at 37°C. Immediately following transposition, the sample was purified by means of a Qiagen MinElute kit.
3. *PCR*. After purification, the library of fragments was amplified using 1 x NEBnext PCR master mix and 1.25µM of custom Nextera PCR primers 1 and 2 (Supplementary Table x). The following PCR conditions were utilized: 72°C for 5 minutes; 98°C for 30 seconds; and thermocycling at 98°C for 10 seconds, 63°C for 30 seconds, and 72°C for 1 minute. In order to reduce GC and size bias in the PCR analysis, PCR reactions were monitored using qPCR in order to stop amplification prior to saturation. This was done by amplification of the full libraries for five cycles, after which an aliquot of the PCR reaction was taken and 10µl of the PCR cocktail containing Sybr Green at a final concentration of 0.6x was added. This reaction was run for 20 cycles in order to determine the additional necessary number of cycles for the remaining 45µl reaction. The libraries were purified by means of a Qiagen PCR cleanup kit that yielded a final library concentration of ∼30nM in 20µl. These libraries were amplified for a total of 10-12 cycles in order to generate the ATAC-seq libraries, which were prepared using the manufacturer’s protocol (Illumina) and paired-end sequenced by means of Illumina NextSeq500 at Cornell University Biotechnology Resource Center.

### ATAC-seq read alignment and peak calling

ATAC-seq sequencing reads were aligned to the mouse reference genome (mm10) using Bowtie2^72^ and filtered for uniquely mapping pairs with a custom python script^73^. Duplicate and multiple aligning reads were removed from the analysis with picardtools v2.1.1 “MarkDuplicates.” Mapped reads were converted to bigWig format using BedTools. To normalize for sequencing depth, libraries were scaled using reads per kilobase sequence per million mapped reads (RPM). ATAC-seq peaks were called on replicate samples using Genrich v0.6 (available at https://github.com/jsh58/Genrich; parameters: -j -f all.log -q 0.05 -a 20.0 -v -e chrM -y). All peaks called by the peak calling software were included in our analysis. Mean FRiP scores for ATAC-seq libraries was 56.7%. The WashU Epigenome Browser was used for visualization of ATAC-seq libraries.

### Principle component analysis

Principle component analysis of the leChRO-seq, RNA-seq, and ATAC-seq datasets were calculated in R v3.4.2^74^ using the *prcomp* function of the R *stats* package^74^. Sequencing reads from each genomics assay were counted in gene bodies and used as input for the PCA analysis. We compared the first two PCs for all datasets.

### Gene transcription analyses

#### Differential expression analysis (DESeq2) of annotated genes for leChRO-seq

We counted reads using the *bigWig* package^75^ in R v3.4.2 in the intervals between 1) the annotated transcription start site (TSS) to 250 bp downstream of the annotated TSS; 2) 250 bp downstream of the annotated TSS to the 3’ end of the gene; 3) the 3’ end of the gene to 5000 bp downstream of the 3’ end of the gene. These windows were selected to count reads within the pause peak, gene body, and post poly(A) signals of all annotated genes. We limited analyses to gene annotations longer than 500 bp in length and gene annotations in which the end of the gene was at least 5000 bp away from the TSS of neighboring genes. Differential expression analysis was conducted using DESeq2^33^ v1.32.0 (parameters: fit type = “local”) and differentially expressed genes were defined as those with a false discovery rate (FDR) less than 0.05. Normalization of the read count matrix was achieved by manually selecting DESeq2 size factors by dividing the read count matrix by library size and multiplying by the relative transcriptional activity of the prophase I substages for each replicate (above). DESeq2 was conducted to compare differential expression between prophase I substages.

#### Scaled metagene plot

We used a publicly available script to generate leChRO metaplots (https://github.com/Danko-Lab/histone-mark-imputation/figures/ScaledMetaPlotFunctions.R). This script plots the average leChRO-seq signal intensity for all annotated genes in Fig. 2. The final plots represent a median of 1000 subsamples, without replacement. Scaled metagene plots were generated using bigWigs normalized to FPKM and relative transcriptional activity for each substage (above).

#### Pausing index and post polyadenylation site (PAS) retention index

Violin plot (Fig. 2) and box and whisker plots (Fig. 3) of the pausing index from the leChRO-seq libraries were calculated in R v3.4.2 by determining the ratio of the leChRO-seq density at the 5’ end of the gene to that in the gene body. Box and whisker plots of the post PAS retention index for clusters 1 and 2, presented in Fig. 3, were calculated in R v.4.0.5 by determining the ratio of the leChRO-seq density in the gene body to that in the 3’ end of the gene to 5000 bp downstream. Statistical analyses performed used the Wilcoxon matched-pairs signed rank test. Alpha values were established at 0.05.

#### MA plots

MA plots of the leChRO-seq datasets were calculated in R v 4.0.5 using the *plotMA* function of the *DESeq2* package.

#### Metaplots

Metaplots in Fig. 5 show the average signal of the sites being summarized using the *metaplot*.*bigWig* function of the *bigWig* R package. leChRO-seq signal was compared to publicly available mapped SPO11 oligos^47^, PRDM9 binding sites^47^, and DMC1 binding sites^51^.

#### Clustering analysis and heatmap

The Python package *clust*^*76*^ v1.12.0 was used to run *clust* (parameters: -n 3 -t 1.0) on the normalized read count matrix for all gene intervals of differentially expressed genes (above). The read count matrix for P and D were normalized to the read count for LZ. The *pheatmap* package^77^ in R v4.0.5 was used to generate the heatmap in Fig. 3.

#### dREG

dREG was run using the default settings to identify putative transcriptional regulatory elements (TREs) from leChRO-seq libraries^28^. A complete description of dREG can be found here: https://github.com/Danko-Lab/dREG.

#### Transcription factor binding site motif analysis for dREG peaks

Briefly, we merged dREG peaks from all prophase I stages using BedTools “merge” and counted leChRO-seq reads for each prophase I substage using the *bigWig* package in R v3.4.2. Differential expression of dREG peaks comparing LZ to P and P to D was performed using DESeq2 (for normalization scheme, see above). Volcano plots reflecting DE of dREG peaks were made using the *EnhancedVolcano* package^78^ v1.10.0 in R. dREG peaks with a log_2_FC > 1 between LZ and P were selected to identify the top DE dREG peaks (28,232 total peaks) for transcription factor (TF) binding motif enrichment. HOMER^39^ was used to identify enriched TF binding motifs within upregulated dREG peaks between LZ and P, allowing multiple motifs per peak and searching for up to 25 motifs 100 bp upstream and downstream from the center of each peak. Donut plot of enriched TF families was generated using the *lessR* package^79^ v4.0.5 in R. MotifScan was used to determine the genomic position of upregulated TREs in P containing the A-MYB binding motif^80^. BedTools “closest” was used to identify putative DE target genes with an FDR < 0.05 for each TRE containing the A-MYB binding motif. Putative DE target genes were defined as those nearest to each TRE containing the A-MYB motif within 50 kb.

#### GO annotation and enrichment

The GO enrichment analysis shown in Figs. 3 and 4 were carried out using the R package *clusterProfiler*^*37*^ v4.0.5. In Fig. 3, subsets of genes identified by DE analysis carried out between prophase I substages (above) were selected based on clustering results. In Fig. 4, DEG within 50 kb of TREs were selected for the A-MYB transcription factor binding motif. These genes were used as input to the enrichGO function. The Benjamini-Hochberg method (a = 0.05) was applied to control for FDR. GO annotation enrichment for biological processes found in Fig. 3 were graphically represented using the *dotplot* function of the *Enrichplot*^*81*^ v1.12.2 package. Dot plots were ranked by adjusted *p*-value and gene set size. GO annotation enrichment for biological processes found in Fig. 4 were made using the *cnetplot* function of the *Enrichplot* package.

#### A-MYB and BRDT ChIP-seq data analysis

Previously published datasets for A-MYB ChIP-seq peaks^20^ and BRDT ChIP-seq peaks^44^ were remapped from the mm9 to mm10 reference genome and mm8 to mm10 reference genome, respectively, using the UCSC LiftOver tool. Bar plots of the percent of A-MYB or BRDT peaks overlapping with the TSSs of genes in cluster 1, cluster 2, or all transcription units (TUs) (Fig. 4C-D) were generated using BedTools intersect and plotted with GraphPad Prism. Statistical analyses comparing the percent of A-MYB or BRDT peaks overlapping with the TSSs of genes in cluster 1 and cluster 2 used Fisher’s exact test. Statistical analyses comparing the percent of A-MYB or BRDT peaks overlapping with the TSSs of genes in cluster 1 or cluster 2 and all TUs used the Chi-squared test with Yates’ correction. Bar plots showing the min, max, and median pausing index (Fig. 4F-I) from leChRO-seq libraries for genes bound by A-MYB or BRDT or not bound by A-MYB or BRDT were calculated in R v4.0.5 by determining the ratio of the leChRO-seq density at the 5’ end of the gene to that in the gene body and plotted using GraphPad Prism. Statistical analyses performed used the Wilcoxon matched-pairs signed rank test. All alpha values were established at 0.05. Metaplots and heatmaps of ATAC-seq signal centered on A-MYB ChIP-seq peaks were generated with deepTools^20,82^.

#### RNA-seq analysis

Quantification of sequencing reads was performed using Salmon^83^ v1.5.2 to the mm10 genome and *tximport*^*84*^ v1.20.0 was used to summarize transcript-level to gene-level abundance estimates in R v4.0.5. Raw sequencing counts from Salmon were DESeq2 normalized (parameters: rowSums(counts(dds)) > 1). DE genes were defined as those with an FDR less than 0.05. Scatterplots comparing the log_2_ fold change and log FPKM of DE genes or all genes using leChRO-seq and RNA-seq data as input presented in Fig. S4 were generated with *ggplot2*^*85*^.

#### Analysis of ATAC-seq peaks

Peak calls from biological replicates for each prophase I substage were used to determine total unique peak sets for each substage. Peak calls representing all of prophase I were then combined and merged using BedTools. Both transcribed and non-transcribed ATAC-seq peaks were then compared to a publicly available H3K4me3 dataset^86^ and annotated TSSs using BedTools intersect. Counts of reads falling into merged peak files were determined using the BedTools multicov function for each sample, and were subsequently RPKM normalized. Mean values + 95% confidence intervals of RPKM for each class of ATAC-seq peaks were presented in **Fig. 5** and plotted using GraphPad Prism. Statistical analyses performed used the Wilcoxon matched-pairs signed rank test. Alpha values were established at 0.05. Metaplots and heatmaps of ATAC-seq signal and H3K4me3 signal centered on SPO11 oligos were generated with deepTools^47,82,86^.

#### Analysis of overlap of accessibility, transcription initiation, and DSB markers

We first asked whether transcription initiation, marked by dREG peaks, and the DSB markers (PRDM9 binding motifs, Spo11 oligos, and DMC1 ChIP-seq peaks) are enriched in accessible chromatin, based on overlap with ATAC-seq peaks at the different stages of meiotic prophase I^47,51^. Using the center of the dREG or the DSB marker peaks, we asked what percentage of the ATAC-seq peaks overlap with either of these markers by counting the number of ATAC-seq peaks with values greater than zero for any of the markers following BedTools “intersect” with -C function, at all three stages. To ask if the calculated overlaps between ATAC-seq peaks and the different markers are different than what is expected by chance, we randomly shuffled the positions of ATAC-seq peaks across the genome, while preserving their size, preventing overlaps or changes in chromosome distribution with BedTools “shuffle -chrom -noOverlapping”, 1000 times. In each of these 1000 iterations we asked what percentage of the shuffled ATAC-seq peaks overlapped with the dREG and DSB markers. We then calculated an empirical p-value for each marker by taking the fraction of the iterations that resulted in ATAC-seq peak overlaps greater than the observed overlap.

We then asked if the accessible genomic regions presenting transcription initiation occur near or are separated from sites of DSBs. We considered a window of 5kb around the center of ATAC-seq peaks in each of the stages and asked what fraction of these expanded peaks overlapped with each of the markers, including dREG and DSB markers. We then calculated the co-occurrence proportion of dREG and DSB markers in the same expanded ATAC-seq peak. We defined the column of dREG overlapping expanded ATAC-seq peaks as “transcribed ATAC peaks” and randomly shuffled this column 1000 times while reassessing the co-occurrence in each iteration. The empirical p-value was calculated by the number of iterations in which the number of ATAC-seq peaks overlapping with dREG (shuffled “transcribed ATAC peaks”) and the DSB markers was smaller than the observed co-occurrence calculated.

#### BioRender

Figures 2A and 4J were created with BioRender.com.

## ACKNOWLEDGEMENTS

We thank J. Lewis and C. Pereira for sharing scripts used in data analysis, J. Lis and A. Ozer for valuable discussions on radioactive run-ons, and J. Grenier and the TREx core for preparing RNA-seq libraries. Valuable discussions with A. Grimson, J. Schimenti, M. Roberson, and all members of the Danko and Cohen labs during the course of this project helped to shape our manuscript. Work in this publication was supported by R01-HG009309 (NHGRI) to CGD and P50-HD104454 (NICHD) to PEC and CGD. The content is solely the responsibility of the authors and does not necessarily represent the official views of the US National Institutes of Health.

## REFERENCES

1. Griswold, M. D. Spermatogenesis: the commitment to meiosis. Physiol. Rev. 96, 1–17 (2016).

2. Gray, S. & Cohen, P. E. Control of meiotic crossovers: from double-strand break formation to designation. Annu. Rev. Genet. 50, 175–210 (2016).

3. Keeney, S. Spo11 and the Formation of DNA Double-Strand Breaks in Meiosis. Genome Dyn. Stab. 2, 81–123 (2008).

4. Baudat, F., Imai, Y. & de Massy, B. Meiotic recombination in mammals: localization and regulation. Nat. Rev. Genet. 14, 794–806 (2013).

5. Page, S. L. & Hawley, R. S. The genetics and molecular biology of the synaptonemal complex. Annu. Rev. Cell Dev. Biol. 20, 525–558 (2004).

6. Xu, H. et al. Molecular organization of mammalian meiotic chromosome axis revealed by expansion STORM microscopy. Proc Natl Acad Sci USA 116, 18423–18428 (2019).

7. Kauppi, L. et al. Numerical constraints and feedback control of double-strand breaks in mouse meiosis. Genes Dev. 27, 873–886 (2013).

8. Geisinger, A., Rodríguez-Casuriaga, R. & Benavente, R. Transcriptomics of meiosis in the male mouse. Front. Cell Dev. Biol. 9, 626020 (2021).

9. Soumillon, M. et al. Cellular source and mechanisms of high transcriptome complexity in the mammalian testis. Cell Rep. 3, 2179–2190 (2013).

10. Hammoud, S. S. et al. Chromatin and transcription transitions of mammalian adult germline stem cells and spermatogenesis. Cell Stem Cell 15, 239–253 (2014).

11. Luo, Z. et al. Reorganized 3D genome structures support transcriptional regulation in mouse spermatogenesis. iScience 23, 101034 (2020).

12. Maezawa, S., Yukawa, M., Alavattam, K. G., Barski, A. & Namekawa, S. H. Dynamic reorganization of open chromatin underlies diverse transcriptomes during spermatogenesis. Nucleic Acids Res. 46, 593–608 (2018).

13. da Cruz, I. et al. Transcriptome analysis of highly purified mouse spermatogenic cell populations: gene expression signatures switch from meiotic-to postmeiotic-related processes at pachytene stage. BMC Genomics 17, 294 (2016).

14. Grive, K. J. et al. Dynamic transcriptome profiles within spermatogonial and spermatocyte populations during postnatal testis maturation revealed by single-cell sequencing. PLoS Genet. 15, e1007810 (2019).

15. Naro, C. et al. An Orchestrated Intron Retention Program in Meiosis Controls Timely Usage of Transcripts during Germ Cell Differentiation. Dev. Cell 41, 82–93.e4 (2017).

16. Royo, H. et al. Evidence that meiotic sex chromosome inactivation is essential for male fertility. Curr. Biol. 20, 2117–2123 (2010).

17. Manterola, M. et al. BRDT is an essential epigenetic regulator for proper chromatin organization, silencing of sex chromosomes and crossover formation in male meiosis. PLoS Genet. 14, e1007209 (2018).

18. Matzuk, M. M. et al. Small-molecule inhibition of BRDT for male contraception. Cell 150, 673–684 (2012).

19. Bolcun-Filas, E. et al. A-MYB (MYBL1) transcription factor is a master regulator of male meiosis. Development 138, 3319–3330 (2011).

20. Li, X. Z. et al. An ancient transcription factor initiates the burst of piRNA production during early meiosis in mouse testes. Mol. Cell 50, 67–81 (2013).

21. Maezawa, S. et al. Super-enhancer switching drives a burst in gene expression at the mitosis-to-meiosis transition. Nat. Struct. Mol. Biol. 27, 978–988 (2020).

22. Özata, D. M. et al. Evolutionarily conserved pachytene piRNA loci are highly divergent among modern humans. Nat. Ecol. Evol. 4, 156–168 (2020).

23. Chu, T. et al. Chromatin run-on and sequencing maps the transcriptional regulatory landscape of glioblastoma multiforme. Nat. Genet. 50, 1553–1564 (2018).

24. Abe, H. et al. The Initiation of Meiotic Sex Chromosome Inactivation Sequesters DNA Damage Signaling from Autosomes in Mouse Spermatogenesis. Curr. Biol. 30, 408–420.e5 (2020).

25. Noe Gonzalez, M., Blears, D. & Svejstrup, J. Q. Causes and consequences of RNA polymerase II stalling during transcript elongation. Nat. Rev. Mol. Cell Biol. 22, 3–21 (2021).

26. Rougvie, A. E. & Lis, J. T. The RNA polymerase II molecule at the 5’ end of the uninduced hsp70 gene of D. melanogaster is transcriptionally engaged. Cell 54, 795–804 (1988).

27. Love, J. D. & Minton, K. W. Screening of λ library for differentially expressed genes using in vitro transcripts. Anal. Biochem. 150, 429–441 (1985).

28. Wang, Z., Chu, T., Choate, L. A. & Danko, C. G. Identification of regulatory elements from nascent transcription using dREG. Genome Res. 29, 293–303 (2019).

29. Widger, A. et al. ATR is a multifunctional regulator of male mouse meiosis. Nat. Commun. 9, 2621 (2018).

30. Mullen, T. E. & Marzluff, W. F. Degradation of histone mRNA requires oligouridylation followed by decapping and simultaneous degradation of the mRNA both 5’ to 3’ and 3’ to 5’. Genes Dev. 22, 50–65 (2008).

31. National Center for Biotechnology Information, U.S. National Library of Medicine. 1700065D16Rik RIKEN cDNA 1700065D16 gene [Mus musculus (house mouse)]. https://www.ncbi.nlm.nih.gov/gene/73410 (2021).

32. Siepel, A. A unified probabilistic modeling framework for eukaryotic transcription based on nascent RNA sequencing data. BioRxiv (2021) doi:10.1101/2021.01.12.426408.

33. Love, M. I., Huber, W. & Anders, S. Moderated estimation of fold change and dispersion for RNA-seq data with DESeq2. Genome Biol. 15, 550 (2014).

34. Yang, F., Skaletsky, H. & Wang, P. J. Ubl4b, an X-derived retrogene, is specifically expressed in post-meiotic germ cells in mammals. Gene Expr. Patterns 7, 131–136 (2007).

35. Shalini, V., Bhaduri, U., Ravikkumar, A. C., Rengarajan, A. & Satyanarayana, R. M. R. Genome-wide occupancy reveals the localization of H1T2 (H1fnt) to repeat regions and a subset of transcriptionally active chromatin domains in rat spermatids. Epigenetics Chromatin 14, 3 (2021).

36. Jha, K. N., Tripurani, S. K. & Johnson, G. R. TSSK6 is required for γH2AX formation and the histone-to-protamine transition during spermiogenesis. J. Cell Sci. 130, 1835–1844 (2017).

37. Wu, T. et al. clusterProfiler 4.0: A universal enrichment tool for interpreting omics data. Innovation (Camb) 2, 100141 (2021).

38. Best, D. J. & Roberts, D. E. Algorithm AS 89: the upper tail probabilities of spearman’s rho. Appl. Stat. 24, 377 (1975).

39. Heinz, S. et al. Simple combinations of lineage-determining transcription factors prime cis-regulatory elements required for macrophage and B cell identities. Mol. Cell 38, 576–589 (2010).

40. Horvath, G. C., Kistler, M. K. & Kistler, W. S. RFX2 is a candidate downstream amplifier of A-MYB regulation in mouse spermatogenesis. BMC Dev. Biol. 9, 63 (2009).

41. Don, J. & Stelzer, G. The expanding family of CREB/CREM transcription factors that are involved with spermatogenesis. Mol. Cell. Endocrinol. 187, 115–124 (2002).

42. Wu, Y. et al. Transcription factor RFX2 is a key regulator of mouse spermiogenesis. Sci. Rep. 6, 20435 (2016).

43. Cheung, K. L., Kim, C. & Zhou, M.-M. The functions of BET proteins in gene transcription of biology and diseases. Front. Mol. Biosci. 8, 728777 (2021).

44. Her, Y. R. et al. Genome-wide chromatin occupancy of BRDT and gene expression analysis suggest transcriptional partners and specific epigenetic landscapes that regulate gene expression during spermatogenesis. Mol. Reprod. Dev. 88, 141–157 (2021).

45. Baudat, F. et al. PRDM9 is a major determinant of meiotic recombination hotspots in humans and mice. Science 327, 836–840 (2010).

46. Parvanov, E. D., Petkov, P. M. & Paigen, K. Prdm9 controls activation of mammalian recombination hotspots. Science 327, 835 (2010).

47. Lange, J. et al. The Landscape of Mouse Meiotic Double-Strand Break Formation, Processing, and Repair. Cell 167, 695–708.e16 (2016).

48. Grey, C., Baudat, F. & de Massy, B. PRDM9, a driver of the genetic map. PLoS Genet. 14, e1007479 (2018).

49. Diagouraga, B. et al. PRDM9 Methyltransferase Activity Is Essential for Meiotic DNA Double-Strand Break Formation at Its Binding Sites. Mol. Cell 69, 853–865.e6 (2018).

50. Patel, L. et al. Dynamic reorganization of the genome shapes the recombination landscape in meiotic prophase. Nat. Struct. Mol. Biol. 26, 164–174 (2019).

51. Brick, K., Smagulova, F., Khil, P., Camerini-Otero, R. D. & Petukhova, G. V. Genetic recombination is directed away from functional genomic elements in mice. Nature 485, 642–645 (2012).

52. Zuo, W. et al. Stage-resolved Hi-C analyses reveal meiotic chromosome organizational features influencing homolog alignment. Nat. Commun. 12, 5827 (2021).

53. Vara, C. et al. Three-Dimensional Genomic Structure and Cohesin Occupancy Correlate with Transcriptional Activity during Spermatogenesis. Cell Rep. 28, 352–367.e9 (2019).

54. Margolin, G., Khil, P. P., Kim, J., Bellani, M. A. & Camerini-Otero, R. D. Integrated transcriptome analysis of mouse spermatogenesis. BMC Genomics 15, 39 (2014).

55. Gilchrist, D. A. et al. Pausing of RNA polymerase II disrupts DNA-specified nucleosome organization to enable precise gene regulation. Cell 143, 540–551 (2010).

56. Cohen, P. E., Pollack, S. E. & Pollard, J. W. Genetic analysis of chromosome pairing, recombination, and cell cycle control during first meiotic prophase in mammals. Endocr. Rev. 27, 398–426 (2006).

57. Walker, M. et al. Affinity-seq detects genome-wide PRDM9 binding sites and reveals the impact of prior chromatin modifications on mammalian recombination hotspot usage. Epigenetics Chromatin 8, 31 (2015).

58. Holloway, J. K., Sun, X., Yokoo, R., Villeneuve, A. M. & Cohen, P. E. Mammalian CNTD1 is critical for meiotic crossover maturation and deselection of excess precrossover sites. J. Cell Biol. 205, 633–641 (2014).

59. Gray, S., Santiago, E. R., Chappie, J. S. & Cohen, P. E. Cyclin N-Terminal Domain-Containing-1 Coordinates Meiotic Crossover Formation with Cell-Cycle Progression in a Cyclin-Independent Manner. Cell Rep. 32, 107858 (2020).

60. Kolas, N. K. et al. Localization of MMR proteins on meiotic chromosomes in mice indicates distinct functions during prophase I. J. Cell Biol. 171, 447–458 (2005).

61. Bryant, J. M., Meyer-Ficca, M. L., Dang, V. M., Berger, S. L. & Meyer, R. G. Separation of spermatogenic cell types using STA-PUT velocity sedimentation. J. Vis. Exp. (2013) doi:10.3791/50648.

62. Gaysinskaya, V., Soh, I. Y., van der Heijden, G. W. & Bortvin, A. Optimized flow cytometry isolation of murine spermatocytes. Cytometry A 85, 556–565 (2014).

63. Mahat, D. B. et al. Base-pair-resolution genome-wide mapping of active RNA polymerases using precision nuclear run-on (PRO-seq). Nat. Protoc. 11, 1455–1476 (2016).

64. Schmieder, R. & Edwards, R. Quality control and preprocessing of metagenomic datasets. Bioinformatics 27, 863–864 (2011).

65. Martin, M. sequencer. Other solutions for adapter trimming exist. Some software libraries, such as HTSeq by Simon Anders (.

66. Li, H. & Durbin, R. Fast and accurate short read alignment with Burrows-Wheeler transform. Bioinformatics 25, 1754–1760 (2009).

67. Quinlan, A. R. & Hall, I. M. BEDTools: a flexible suite of utilities for comparing genomic features. Bioinformatics 26, 841–842 (2010).

68. Li, D., Hsu, S., Purushotham, D., Sears, R. L. & Wang, T. WashU Epigenome Browser update 2019. Nucleic Acids Res. 47, W158–W165 (2019).

69. Dobin, A. et al. STAR: ultrafast universal RNA-seq aligner. Bioinformatics 29, 15–21 (2013).

70. Buenrostro, J. D., Giresi, P. G., Zaba, L. C., Chang, H. Y. & Greenleaf, W. J. Transposition of native chromatin for fast and sensitive epigenomic profiling of open chromatin, DNA-binding proteins and nucleosome position. Nat. Methods 10, 1213–1218 (2013).

71. Buenrostro, J. D., Wu, B., Chang, H. Y. & Greenleaf, W. J. ATAC-seq: a method for assaying chromatin accessibility genome-wide. Curr. Protoc. Mol. Biol. 109, 21.29.1-21.29.9 (2015).

72. Langmead, B. & Salzberg, S. L. Fast gapped-read alignment with Bowtie 2. Nat. Methods 9, 357–359 (2012).

73. Lewis, J. J. & Reed, R. D. Genome-Wide Regulatory Adaptation Shapes Population-Level Genomic Landscapes in Heliconius. Mol. Biol. Evol. 36, 159–173 (2019).

74. Team, R. C. R: A language and environment for statistical computing. 2015. R foundation for statistical computation, Vienna, Austria. (2020).

75. Andre L. Martins, copyright holder Cornell University. src/jkweb by Jim Kent, copyright by Jim Kent and Regents of the University of California. (2014). bigWig: R Interface to Query UCSC BigWig Files. R package version 0.2-9.

76. Abu-Jamous, B. & Kelly, S. Clust: automatic extraction of optimal co-expressed gene clusters from gene expression data. Genome Biol. 19, 172 (2018).

77. Kolde, R. & Kolde, M. R. Package ‘pheatmap’. R package 1, (2015).

78. Kevin Blighe, Sharmila Rana and Myles Lewis (2020). EnhancedVolcano: Publication-ready volcano plots with enhanced colouring and labeling. R package version 1.8.0. https://github.com/kevinblighe/EnhancedVolcano.

79. Gerbing, D. W. lessR: Less Code, More Results. (2021).

80. Sun, H. et al. Quantitative integration of epigenomic variation and transcription factor binding using MAmotif toolkit identifies an important role of IRF2 as transcription activator at gene promoters. Cell Discov. 4, 38 (2018).

81. Guangchuang Yu (2021). enrichplot: Visualization of Functional Enrichment Result. R package version 1.10.2. https://yulab-smu.top/biomedical-knowledge-mining-book/.

82. Ramírez, F., Dündar, F., Diehl, S., Grüning, B. A. & Manke, T. deepTools: a flexible platform for exploring deep-sequencing data. Nucleic Acids Res. 42, W187–91 (2014).

83. Patro, R., Duggal, G., Love, M. I., Irizarry, R. A. & Kingsford, C. Salmon provides fast and bias-aware quantification of transcript expression. Nat. Methods 14, 417–419 (2017).

84. Soneson, C., Love, M. I. & Robinson, M. D. Differential analyses for RNA-seq: transcript-level estimates improve gene-level inferences. [version 2; peer review: 2 approved]. F1000Res. 4, 1521 (2015).

85. Wickham, H., Chang, W. & Wickham, M. H. Package ‘ggplot2’. Create Elegant Data Visualisations Using the Grammar of Graphics. Version 2, 1–189 (2016).

86. Maezawa, S. et al. Polycomb protein SCML2 facilitates H3K27me3 to establish bivalent domains in the male germline. Proc Natl Acad Sci USA 115, 4957–4962 (2018).

